# Network shape intelligence outperforms AlphaFold2 intelligence in vanilla protein interaction prediction

**DOI:** 10.1101/2023.08.10.552825

**Authors:** Ilyes Abdelhamid, Alessandro Muscoloni, Danny Marc Rotscher, Matthias Lieber, Ulf Markwardt, Carlo Vittorio Cannistraci

## Abstract

For decades, scientists and engineers have been working to predict protein interactions, and network topology methods have emerged as extensively studied techniques. Recently, approaches based on AlphaFold2 intelligence, exploiting 3D molecular structural information, have been proposed for protein interaction prediction, they are promising as potential alternatives to traditional laboratory experiments, and their design and performance evaluation is compelling.

Here, we introduce a new concept of intelligence termed Network Shape Intelligence (NSI). NSI is modelled via network automata rules which minimize external links in local communities according to a brain-inspired principle, as it draws upon the local topology and plasticity rationales initially devised in brain network science and then extended to any complex network. We show that by using only local network information and without the need for training, these network automata designed for modelling and predicting network connectivity can outperform AlphaFold2 intelligence in vanilla protein interactions prediction. We find that the set of interactions mispredicted by AlphaFold2 predominantly consists of proteins whose amino acids exhibit higher probability of being associated with intrinsically disordered regions. Finally, we suggest that the future advancements in AlphaFold intelligence could integrate principles of NSI to further enhance the modelling and structural prediction of protein interactions.

## Introduction

AlphaFold2^1^ (AF2) is a deep-learning artificial intelligence (AI) algorithm that has made accurate the computational prediction of protein structures from amino acid sequences, attaining unimaginable performance in the blind test for protein structure prediction CASP14^1^. Recently, a second type of AI based on large language modelling called ESMFold^2^ has been introduced. ESMFold achieves lower performance than AF2, but it does not require multiple sequence alignment (MSA). Meanwhile, Evans et al. have released AlphaFold2-multimer (AFM), a refined version of AlphaFold2 for the prediction of protein complexes^3^. This breakthrough of AF2 and AFM has sparked interest in structural-based (a.k.a. structurally resolved) prediction of protein-protein interactions (PPIs)^4–6^, and its results are viewed as a milestone towards the structurally resolved human protein interaction network^7^. Gao et al.^6^ has shown that the likelihood of complex formation can be assessed by two measures: the interface-score (IS) and the predicted interface TM-score (piTM), both of which evaluate the confidence of the predicted protein-protein interface if found in an assessed model. Each of these two scores range from 0 to 1, higher score indicates higher confidence and can be used to predict interacting protein pairs using AFM. This is not surprising, as the interface structures of PPIs are not fundamentally different from those that drive protein folding. However, for protein interaction prediction the use of homologous structures as templates is crucial, as AFM often fails to predict the correct structure without them^8^. Meanwhile, in network science, the increasing coverage of the protein interactome^9–13^ has also led to the development of local network topology measures for reliability estimation^14^ and prediction of protein interactions^15–17^. These network-based methods exploit topological path-based patterns of already mapped interactions to predict the likelihood of missing interactions. This is a vanilla protein interaction prediction since 3D molecular structural information is neither used for training nor predicted. Hence, the information required as input for network-based methods is the mere network shape: meaning a binary adjacency matrix indicating the presence or absence of an unweighted and undirected link between each pair of network’s nodes. Therefore, the structural-based interaction prediction can be viewed as a sub-case of vanilla protein interaction prediction that not only predicts the likelihood of the interaction but also the 3D molecular structure.

While most local rules for modeling network connectivity were based on paths of length 2^14,15^ (Fig. 1a), in 2015 our group^18^ proposed to use paths of length 3 (Fig. 1b) in complex bipartite networks. We argued that the accepted notion according to which the common neighbor nodes of a complex network should be associated to the triadic closure principle^16^ is incomplete and misleading, and that the closure rule to define the common neighbors is dependent by the connectivity rationale that is behind the complex network topology^18^. For instance, in bipartite networks (Fig. 1b), the connectivity rationale is not based on similarity between nodes but on their complementarity, such as the interactions in recommender systems^19,20^ that can be only between a user and an item (direct interactions between users or between items are not possible). This implies that in bipartite networks the common neighbor nodes are associated to a quadratic closure principle^18^, which is the closure structure of minimum topological length characterizing the complementarity connectivity rationale. Therefore, according to our theoretical framework, common neighbors are the intermediate nodes traversed by all the paths of length 3 between two seed nodes (Fig 1b). In case of paths of length n, we propose to adopt the general definition: ‘local ring’ closure^21^. In 2018, our group^22^ successfully exploited this path-length 3 rationale for network-based prediction of drug-target interactions, which are biological networks based on bipartite complementarity connectivity. In a preprint (in 2018)^23^ and in the respective peer reviewed publication (in 2019)^16^, Kovács et al. highlighted that a path-length 3 complementary connectivity rationale - similar to the one discovered by our group^18^ for bipartite networks - plays a major role also in the monopartite organization of protein interaction networks, because of 3D-structural and evolutionary bio-principles that govern this class of monopartite networks. They proposed a new path-length 3 measure called L3 for topological prediction of protein interactions, and they showed that L3 could outperform Cannistraci Resource Allocation (CRA)^15^ which was the best performing path-length 2 method in literature out of 23 different methods tested in their study^16^. Kovács et al. L3 network modelling was introduced according to biologically sound rationales valid for protein networks only, and its mathematical definition as a mechanistic model lacks derivation from current theoretical knowledge, missing any connection to already known generalized principles for modelling and prediction of network connectivity.

**Figure 1.**
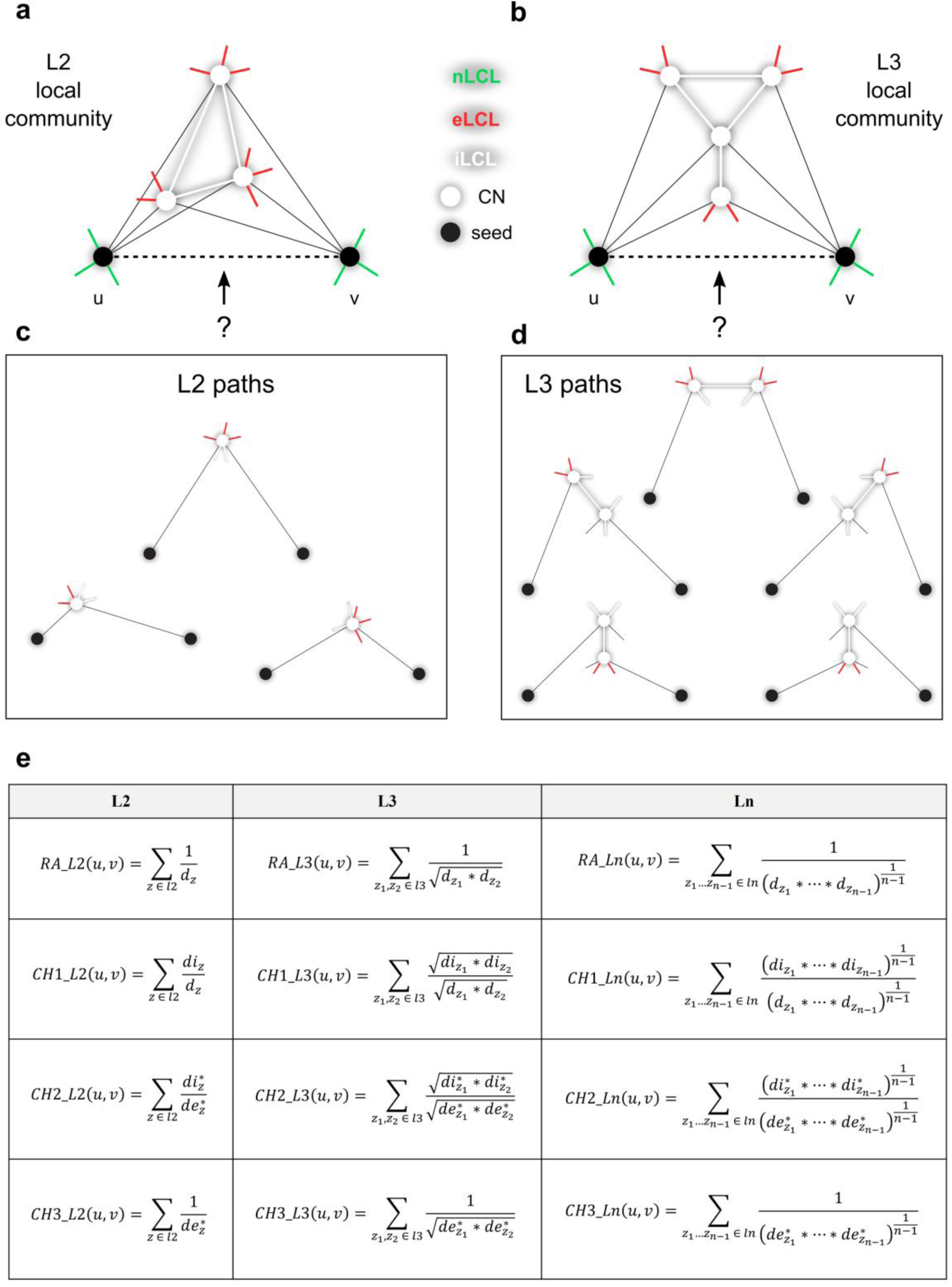
Cannistraci-Hebb epitopological rationale and network automata on paths of length 2, 3 and n. Examples for local topological link prediction performed using the (**a**) L2 or (**b**) L3 Cannistraci-Hebb epitopological rationale. The two black nodes represent the seed nodes whose nonobserved interaction should be scored with a likelihood. The white nodes are the L2 or L3 common-neighbours (CNs) of the seed nodes, further neighbours are not shown for simplicity. The different types of links are reported with different colours: non-LCL (nLCL, green), external-LCL (eLCL, red), internal-LCL (iLCL, white). The cohort of common-neighbours and the iLCL (white links) form the local community only if the external links (red links) are minimized, which is the Cannistraci-Hebb epitopological rationale and the necessary condition to be a Cannistraci-Hebb network automaton. The set of (**c**) L2 and (**d**) L3 paths related to the given examples of local communities are shown. (**e**) Mathematical definitions of Cannistraci-Hebb network automata on path of length 2 (*L2*), 3 (*L3*) and n (*Ln*). Notation: *u*, *v* are the seed nodes; *z* is the intermediate node (CN); *d*_*z*_ is the degree of *z*; *di*_*z*_ is the internal degree (number of iLCL) of *z*; *de*_*z*_ is the external degree (number of eLCL) of *z*; *d*^∗^ = 1 + *d*.

In 2018, around 2 months after and in reply to the pre-print of Kovács et al., our group published a pre-print^24^ with four relevant findings that we confirm and further develop in the present study towards the definition of network shape intelligence: (i) a mathematical proof that L3 (renamed RA-L3) formula is the path-length 3 generalization of a well-known path-length 2 modelling principle in network science (specifically in link prediction) which takes the name of resource allocation (RA) and was proposed in 2009 by Tao Zhou et al.^25^; (ii) the proposal of a general theoretical framework for designing local topological measures based on path-length n, where n indicate an arbitrary length; (iii) the definition of the Cannistraci resource allocation in path-length n (CRA-Ln renamed CH1-Ln) and the proposal of the general theory of Cannistraci-Hebb (CH) network automata with a new measure termed CH2-Ln^24^ (and CH3-Ln^26^ that we proposed in a subsequent version of the pre-print); (iv) computational evidence that CH1-L3, CH2-L3 and CH3-L3^26^ significantly outperform L3 on protein interaction networks because they do not minimize interactions between proteins in local communities (Fig.1b). This last finding, since the appearance in our preprints^24,26^, has been confirmed with tests on hundreds of networks in two comparative surveys^27,28^. Here, we define, validate and compare (in 15 different types of protein interactomes) the Cannistraci-Hebb network automata theory versus L3^16^. Based on these results, we propose the definition of what we name: network shape intelligence, which is the intelligence displayed by any topological network automata to perform valid (significantly more than uniformly random) connectivity predictions without training, but only processing the input knowledge associated to the local topological network organization. Then, we propose a computational framework to compare the performance and understand the differences between the power of cutting-edge AF deep learning intelligence and network shape intelligence. This allows us to define a positive and negative set respectively composed by true and false protein interactions, which are used to validate the performance of the protein interaction computational predictions without the need to have at hand knowledge of the native 3D interaction structure. We find that CH network automata models significantly outperform L3 method for prediction of protein interactions tested in 15 networks of different organisms. Then, on Yeast DIP^29–31^ network, we show that regardless of using path-length 2 or path-length 3, CH models significantly outperform AFM and this result is confirmed by using two different performance evaluation measures. Finally, we show that the set of interactions that is wrongly predicted by AFM is composed by proteins with amino acids that on average present significantly higher likelihood to belong to intrinsically disordered regions (IDR).

## Principles and modelling

### From L3 to Resource allocation on paths of length n

Kovács et al.^16^ disclosed a principle based on paths of length three (L3) for network-based prediction of protein interactions that overcomes any rival method based on paths of length two (L2). The L3 principle was motivated by both 3D-structural and evolutionary arguments related to protein interactions, and the mechanistic model (mathematical formula) for modelling the L3 principle in unweighted and undirected networks is:

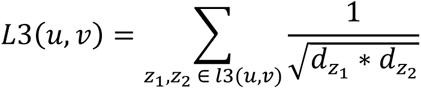

where: *u* and *v* are the two seed nodes of the candidate interaction; *z*_1_ and *z*_2_ are the two intermediate nodes on the considered path of length three *l3(u,v*); *d*_*z*1_ and *d*_*z*2_ are the respective node degrees; and the summation is executed over all the paths of length three. Kovács et al. motivate the penalization for the degree of *z*_1_ and *z*_2_ as follows^16^: << We expect that node pairs connected by the highest number of L3 paths are most likely to be directly connected. However, high degree nodes (hubs) might induce multiple, unspecific shortcuts in the network, biasing the results. To cancel potential degree biases caused by intermediate hubs in the paths, we assign a degree-normalized L3 score to each node pair, *u* and *v* >>.

L3 mathematical definition as a mechanistic model was proposed more as a matter of intuition than as derivation from current theoretical knowledge, and it misses any connection to already known generalized principles for modelling and prediction of network connectivity. *De facto*, this represents a conceptual limitation because it fails to contribute to the theoretical understanding of the universal mechanisms and modelling formalisms that are behind emergence of network connectivity regardless of the specific physical system and its scale.

The first objective of this study is to understand the generalized mechanistic principle of complex network self-organization behind the mathematical formula introduced by Kovács et al.^16^ for modelling the L3 principle in unweighted and undirected networks. And the first important finding is surprisingly simple: the generalized principle behind the L3 formula proposed by Kovács et al.^16^ is already well-known in network science, in particular in link prediction, and is termed *resource allocation* (RA) by Tao Zhou et al.^25^. To prove it, we need a few simple mathematical steps. The basic formula of the RA model on a path of length two (which from here forward we will call RA-L2) is:

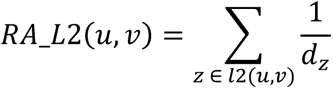

where: *u* and *v* are the two seed nodes of the candidate interaction; *z* is the intermediate node on the considered path of length two *l2(u,v*); *d*_*z*_ is the respective node degree; and the summation is executed over all the paths of length two.

To generalize to paths of length *n* > 2, we need an operator that merges the single contributions of each weighted (in this case degree-penalized) intermediate node on the path of length *n* (common neighbour in the L2 case). If, without loss of generality, we use as merging operator the geometric mean (which is a choice for designing a robust estimator, since if only one intermediate node on the path has a low weight, then the whole path is penalized), we derive the following generalized formula for paths of length *n*:

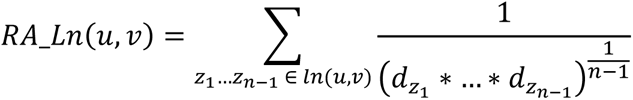

where: *u* and *v* are the two seed nodes of the candidate interaction; *z*_1_… *z*_*n*−1_ are the intermediate nodes on the considered path of length *n ln(u,v*); *d*_*z*1_… *d*_*zn*−1_ are the respective node degrees; and the summation is executed over all the paths of length *n*. For paths of length *n* = 3, the general formula given above becomes clearly equal to L3, which indeed extends the resource allocation principle to paths of length three, therefore from here forward we will call L3 with the new name of RA-L3.

### Network shape intelligence and Cannistraci-Hebb network automata

We define network shape intelligence, the intelligence displayed by any topological network automata^26^ to perform valid (significantly more than random) connectivity predictions without training, by only processing the input knowledge associated to the local topological network organization. The training is not necessary because the network automaton performs predictions extracting information from the network topology, which can be regarded as an associative memory trained directly from the connectivity dynamics of the complex system. In this section we guide the reader along the logical journey that evolves the local-community-paradigm (LCP) theory^14,15,18,21,22,24,26,28^ for link prediction (which is restricted to paths of length 2 or 3), towards the proposed Cannistraci-Hebb theory for modelling network connectivity by means of network automata on paths of length *n*, and ultimately towards the foundation of *network shape intelligence*. To implement this plan, we start by recalling the basic rationale behind the LCP theory, which will turn to be useful also to understand the reasons to introduce the Cannistraci-Hebb theory.

In 1949, Donald Olding Hebb advanced a *local learning rule* in neuronal networks that can be summarized in the following: neurons that fire together wire together^32^. In practice, the Hebbian learning theory assumes that different engrams (memory traces) are memorized by the differing neurons’ cohorts that are co-activated within a given network. Yet, the concept of wiring together was not further specified, and could be interpreted in two different ways. The first interpretation is that the connectivity already present, between neurons that fire together, is reinforced; whereas the second interpretation is the emergence and formation of new connectivity between non-interacting neurons already embedded in an interacting cohort.

In 2013 Cannistraci et al.^15^ noticed that, considering only the network topology, the second interpretation of the Hebbian learning could be formalized as a mere problem of topological link prediction in complex networks. The rationale is the following. The network topology plays a crucial role in isolating cohorts of neurons in functional local communities that allows the neurons to perform local processing. The local-community organization increases the likelihood that a cohort of neurons fires together because they are confined in the same local community, consequently also the likelihood that they will create new connections inside the community is increased by the mere structure of the network topology. This mechanism implements a type of local topological learning that Cannistraci et al.^15^ called *epitopological learning*. *Epitopological learning* occurs when cohorts of neurons tend to be preferentially co-activated, because they are topologically restricted in a local community, and they facilitate learning by forming new connections. Cannistraci et al.^15^ proposed that the identification of this form of learning in neuronal networks was only a special case, hence *epitopological learning* and the associated *local-community-paradigm (LCP)* were proposed as a theory for local learning, self-organization and link-growth valid in general for topological link prediction in any complex network with LCP architecture ^15^. On the basis of these ideas, they proposed a new class of link predictors that demonstrated - also in following studies of other authors^16,27,33–39^ - to outperform many states of the art local-based link predictors both in brain connectomes and in other types of complex networks (such as social, biological, economical, etc.)^15,18,21,22,24^. Furthermore, LCP and epitopological learning can enhance our understanding of how local brain connectivity is able to process, learn and memorize perceptions such as chronic pain ^40^. In conclusion, LCP originated from the initial idea to explain how the network topology indirectly influences the process of learning a memory by adding new connections in a network of neurons, and consequently it was generalized to advocate mechanistic modelling of topological growth and self-organization in real complex networks. How can this theory be exploited also in the domain of prediction of protein interactions?

Protein interactomes display a clear LCP architecture^14,15^, where protein complexes are confined in local and topologically isolated network structures, which are often coincident with functional network modules that play a crucial role in molecular circuits. The key generalized idea behind the LCP network architecture is that, for instance, a local community of neurons or proteins should take functional advantage of being confined in a local assembly of operational units. Each local assembly - if it is properly activated by an external signal coming from another region of the network - performs a functional operation by means of a structural remodelling of the internal connectivity (named iLCL in Fig. 1) between the operational units that are embedded in the network local community. The systems supported by LCP network architecture are very dynamic and react to a stimulus with a local plastic remodelling. In case of operational units such as neurons, the local-community remodelling can implement for instance a learning process. Instead, in case of operational units such as proteins, the local-community remodelling is necessary to implement for instance a biological process, which emerges by the molecular-complex rearrangement in the 3D space. The previous conceptual and mathematical formalizations of the LCP-theory were immature and put more emphasis on the fact that the information content related with the common neighbour nodes should be complemented with the topological information emerging from the interactions between them (the iLCL in Fig. 1). This represents a current limitation of the LCP theory that, by proposing the new concept of Cannistraci-Hebb rule, we want to overcome. Indeed, in 2018 Cannistraci^14^ showed that the local isolation of the operational units in the different local communities is equally important to carve the LCP architecture in the network, and this is guaranteed by the fact that the common neighbours minimize their interactions external (the eLCL in Fig. 1) to the local community^14^. The topological minimization forms a sort of physical and structural ‘energy barrier’, which in turn confines the signal processing to remain internally to the local community^14^. Hereafter, we will revise the LCP idea and its mathematical formalization to explicitly consider also the *minimization of the external* links (the eLCL in Fig. 1). The explicit rule that suggests the minimization of external links is named Cannistraci-Hebb (CH) rule^19,24,26–28^, and all the network automata models that implement in their mathematical formula the CH rule are belonging to the class of CH network automata models^19,21,24,26–28^.

CRA-L2^15^ is the best performing rule in the framework of LCP theory, it represents the LCP extension of RA-L2, and it is a local-ring network automaton model^21^ that seems related and able to predict the growth of network topologies associated to hyperbolic geometry^21^. The starting point to revise the LCP theory is that empirical evidences provided by several studies^16,27,33–39^ in link prediction, and confirmed also by Kovács et al.^16^, show that CRA-L2 outperforms both RA-L2 and the large majority (Kovács et al. tested 23 methods) of other neighbourhood-based mechanistic and parameter-free L2-models. Hence, a fundamental question to address is what mechanisms make CRA-L2 so effective to outperform all the other local-based methods in -length 2 and, most importantly, whether also CRA-L3 would outperform RA-L3. For CRA-L2 network automaton model the likelihood of a new link to appear is function not only of the number of common neighbours, but also function of the numbers of iLCLs and eLCLs (Fig. 1). Therefore, the mathematical formula of CRA in L2 is:

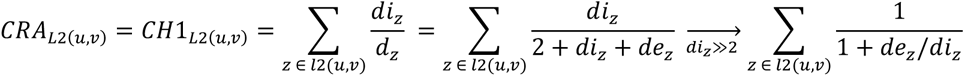

where: *u* and *v* are the two seed nodes of the candidate interaction; *z* is the intermediate node (common neighbour) on the considered path of length two *l2(u,v*); *d*_*z*_ is the respective node degree (which is equal to 2+#iLCLs+#eLCLs, where # stays for number); *di*_*z*_ is the respective internal node degree (number of iLCL); and the summation is executed over all the paths of length two.

However, observing this formula, it emerges that the principle behind CRA-L2 is not a resource allocation penalization. On the contrary, the model is based on the common neighbours’ rewards of the internal links (iLCL) balanced by the penalization of the external links. Hence, the current name is misleading. Since this model is a generalization and reinterpretation of a Hebbian learning local-rule to create new topology in networks proposed by Cannistraci (2018)^14^, we decide to rename CRA as L2-based Cannistraci-Hebb network automaton model number one (CH1-L2), rather than Cannistraci-Resource-Allocation (CRA), which was the name adopted in the previous articles. Yet, the formula of CH1-L2 is conceptually and mathematically awkward. If we want to design a model that is based on the minimization of the eLCL and the maximization of the iLCL, why not to write it directly and explicitly? This is what we propose to realize with the definition of a model that we name CH2-L2 and that theoretically should be a straightforward mathematical formalization to express the principle or rule of topological self-organization behind the Cannistraci-Hebb network automaton modelling. The formula of CH2-L2 is:

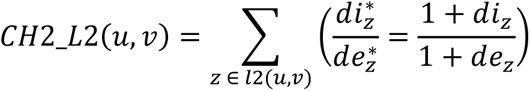

where: *u* and *v* are the two seed nodes of the candidate interaction; *z* is the intermediate node (common neighbour) on the considered path of length two *l2(u,v*); *di*_*z*_ is the respective internal node degree (number of iLCL); *de*_*z*_ is the respective external node degree (number of eLCL); and the summation is executed over all the paths of length two. Note that a unitary term is added to the numerator and denominator to avoid the saturation of the value in case of iLCL or eLCL equal to zero.

To confirm the importance of the isolation of the local communities^14^, we propose CH3-L2, which is a third network automata model that implements exclusively the CH rule, hence its mathematical formula relies on the solely minimization of *de*_*z*_ which is the number of eLCLs for each common neighbour *z* on *l2(u,v)* paths:

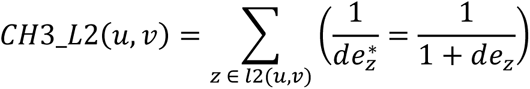

Fig. 1 suggests an interpretation about how the CH network automata models work in a monopartite topology. The ensemble of paths of a given length n creates a local-tunnel which provides a route of connectivity between the two nonadjacent nodes. The higher the number of common neighbours, the higher the size of the local-tunnel. For each common neighbour, the lower the number of eLCLs, the more the shape of the local-tunnel is well-defined and its existence confirmed. Therefore, in link prediction, CH models estimate a likelihood that is proportional both to the size of the local-tunnel (which is associated to the number of common neighbours) and to the extent to which the local-tunnel exists (which is associated to the minimization of external links). In contrast, the classical common neighbour network automaton model only estimates a likelihood proportional to the size of the local-tunnel, which is a significant limitation. Furthermore, the local community represents the central chunk of the local-tunnel exclusively formed by CNs and iLCLs. The local community represents a bridging structure which is fundamental to transfer the information between the two seed nodes. While eLCLs minimization is a necessary condition for the existence of the local-tunnel, the maximization of iLCLs might not play always an important role in all complex systems, and the next section on the experimental evidences will help to shed light on this diversity of the two mechanisms. As a sanity check, Suppl. Table 1 reports the formula of a network automaton that implements the exclusive maximization of iLCLs. The final step in our journey towards the complete definition of CH network automata is the generalization in L*n*. In Suppl. Note 1 and Supp. Table 1, we explain the steps for this generalization and in Fig. 1 we provide the mathematical formula of the CH rules in path-length n.

To summarize, in this study we compare four CH network automata models (RA-Ln, CH1-Ln, CH2-Ln and CH3-Ln) and one not-CH network automaton model (iLCL-Ln) considering path-length 2 and 3. All of these models are based on the definition of local community, but the CH models are the ones that as a necessary condition implement eLCLs minimization, hence note that RA-Ln is also a CH model. As a final remark, we will not consider other similarity measures that are not-CH, such as the many already considered in Kovács et al^16^, because they were already shown to be outperformed by RA-L3. At last, we clarify that the CH model is endowed of a sub-ranking strategy to internally sub-rank all the nonadjacent node pairs characterized by the same CH score. The effect of this sub-ranking strategy is twofold and is discussed in Suppl. Note 1. All the network predictors tested in results section are fairly evaluated considering their adjustment with the sub-ranking strategy.

## Experimental evidences

### Comparing predictions of network shape intelligence methods

Our experiments are based on 10% random removal and re-prediction of the original protein network topology, repeated for 10 realizations: mean and standard error values of performance indicators are considered for the evaluation. This is the standard procedure to test link prediction when - as in protein interactomes - only positive set (observed links in the network) and missing set (the union of negative and undetected positive links) are available^14–19,21–28,41^. Indeed, as we show in Suppl. Fig. 1, the removal of 50% of the original links brings to a significant aberration of the network features, whereas for 10% removal the aberration is negligible. Each predictor assigns a likelihood to the missing links, which are then ranked, and the performance is evaluated using AUP@100^16,24^ which focuses on the precision in the first part (first 100 predicted interactions) of the ranking, AUC-mROC^28,42^ and AUC-PR^43,44^ which are threshold-free measures for unbalanced sets. Indeed, in unbalanced sets, the evaluation purpose is to assess the prediction of few positives in contrast to a large majority of undefined samples, and to assess the extent to which true links are prioritized in all the ranking list^28,42^.

The second important finding of this study (obtained by testing predictions on 15 protein networks) is that CH2-L3 and CH3-L3 models perform significantly better than RA-L3 (Fig. 2, Suppl. Fig. 3,4 and Suppl. Table 2-4), and similarly CH2-L2 and CH3-L2 perform better than RA-L2 (Suppl. Table 2-4). This implies that in general (regardless of L3 or L2 implementation of the model) for protein networks the RA penalization, which was the original principle of self-organization behind the L3 model proposed by Kovács et al., might not be the most appropriate because it minimizes not only the eLCLs (reason for which is a CH model) but also the iLCLs. Hence, in protein interactions, a correct mechanism of prediction might be: i) to adopt L3 paths (as previously discovered by Kovács et al.^16^); ii) to minimize eLCLs (as shown in this study and supported in a previous study of Cannistraci (2018)^14^); iii) to non-minimize iLCLs (as shown in this study and supported in a previous study of Cannistraci (2018)^14^). Indeed, the solely minimization of eLCLs implemented by CH3-L3 seems enough to predict protein interactions using local information, because the iLCL maximization of CH1-L3 and CH2-L3 does not seem to offer significant improvement (Fig. 2, Suppl. Fig. 2-4 and Suppl. Table 2-4). In addition, CH2-L3 and CH3-L3 perform better than CH1-L3 (Suppl. Tables 2-4), and this result indicates that our intuition to design CH2-L3 and CH3-L3, which are more straightforward mathematical formulae for the CH principle, was well-posed.

**Figure 2.**
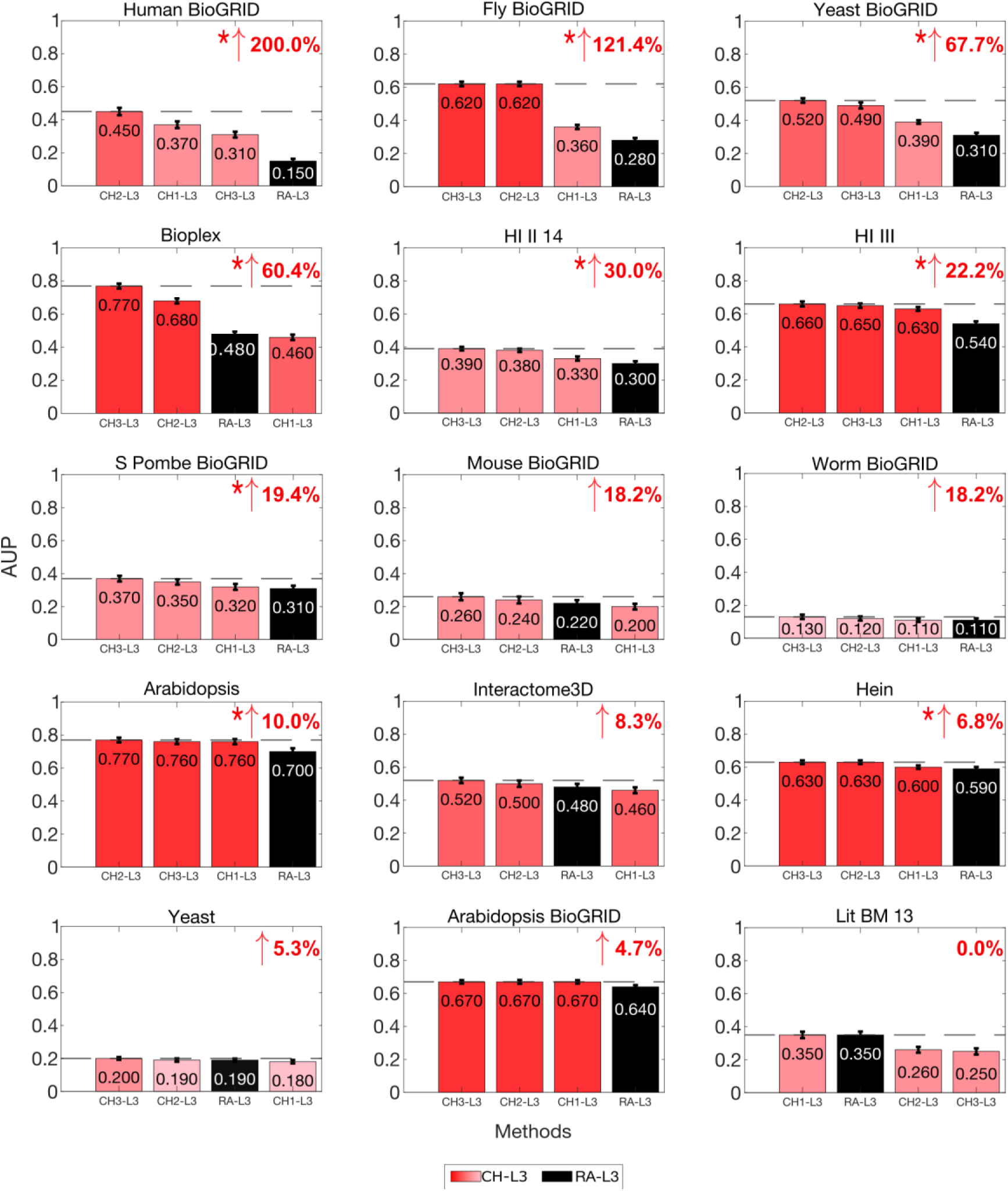
Comparing PPI predictions of network shape intelligence methods. For each network, 10% of links were randomly removed (10 repetitions) and the algorithms were executed to assign likelihood scores to the non-observed links in the reduced networks. To evaluate the performance, the links are ranked by likelihood scores and the precision is computed as the percentage of removed links among the top-r in the ranking, for each r from 1 up to 100 at steps of 1. The barplots show the average (on 10 repetitions) area under precision curve (AUP) and its standard error. The color of the CH bars varies depending on the average AUP value: the higher the average AUP is, the redder the CH bars appear. A red or black arrow is displayed on the top-right of the subplot, indicating the percentage increment between the best CH method and RA-L3. If the arrow is not present, there is not increment. A permutation test for the mean AUP has been conducted and an asterisk next to the arrow indicates significant statistical difference (p-value ≤ 0.05). The networks are ordered (top-down, left-right) by decreasing percentage increment and, in case of tie, by increasing p-value between the two methods.

### Network shape intelligence outperforms AlphaFold2 intelligence in PPI prediction

Since nothing can be concluded about the missing interactions (they are a mix of true and false interactions), the evaluation adopted in the previous section focuses on measuring the extent to which randomly removed positive interactions are prioritized on top of the ranking, because their predicted likelihood is larger than the other missing interactions. However, this procedure is partial, because does not account for the ability of a predictor to put at the bottom of the ranking the interactions that are false (negative). In addition, protein interaction networks have false positive links^14,45^, therefore to evaluate the performance of a predictor we must make sure that the tested positive and negative interactions are reliable. Here, we introduce a computational framework in protein interaction prediction evaluation that tries to overcome these limitations, and we employ it to compare the performance and understand the differences between the power of cutting-edge AF deep learning intelligence and network shape intelligence. Considering the high computational time of AFM for predicting a protein interaction pair, we could focus only on the network of a single organism. Without loss of generality and to ensure that the golden standard of tested interactions is of the highest reliability, we focus on the yeast DIP protein interactome (S. Cerevisiae, DIP database^29–31^, 4951 proteins and 22382 interactions) which is a valuable resource in the field of computational biology, because yeast is one of the most studied organisms^46,47^ and its interactome is one of the most extensively characterized experimentally^11,48^. We propose a bona fide evaluation methodology (BFEM) that can assess the performance in predicting protein interactions using as benchmark the network of any well studied organism such as S. Cerevisiae, from which a golden standard of true and false interactions is generated. The positive set is composed by all existing protein interactions. The golden standard positive (GSP) set is composed by the top 20% positive interactions with the highest value of a gene ontology (GO) semantic measure designed and adopted by Cannistraci (2018) in a previous study on protein interaction reliability^14^. The negative set is composed by selecting an amount of missing interactions that equals the number of existing interactions in the positive set, and with the lowest value of a GO semantic measure designed in this study for this purpose (see Methods). The golden standard negative (GSN) set is composed by the bottom 20% negative interactions with the lowest value of the same GO semantic measure. BFEM applied on the yeast protein network produces a balanced classification scenario with golden standard positive (GSP = 4476) and negative (GSN = 4476) sets. The GSP interactions can be randomly sampled and removed from the network for re-prediction by CH methods; whereas the GSN are not in the network hence, after removal of the GSP, can be directly predicted by CH methods. Contrary to the case of unbalanced prediction scenario discussed in the previous section, here the standard AUC-ROC can be used as performance evaluation measure, together with AUC-PR.

At first, to define the interactions to predict, we consider a BFEM in which we sample uniformly at random 10% positive (2238) and 10% negative (2238) interactions, respectively from the GSP and the GSN (therefore 50% of the GSP and GSN sets). We indicate this BFEM setup as 10%GSP+10%GSN. This procedure is repeated for 10 realizations and the mean prediction performance of the best CH methods (CH2 and CH3 according to the results in previous section) and AFM (AFM-IS and AFM-piTMS^6^) is evaluated. We find that CH3-L3 and CH2-L3 remarkably outperform (Fig. 3, about 40% improvement according to AUC-ROC and 30% improvement according to AUC-PR) AFM-IS and AFM-piTMS, which is the third important finding of our study. The results are confirmed, with even a small improvement of CH methods, if we repeat the analysis considering random 5% positive (1119) and 5% negative (1119) interactions (5%GSP+5%GSN in Suppl. Fig. 5), respectively from the GSP and the GSN. Remarkably, CH3-L3/CH2-L3 outperform AFM-IS/AFM-piTMS of about 70% improvement for AUC-ROC and 60% improvement for AUC-PR when we adopt an evaluation methodology predicting 1% positive (224) and 1% negative (224) interactions (1%positive+1%negative in Suppl. Fig. 6), which are uniformly sampled at random from the entire positive set (all the 22382 observed interactions in the yeast network) and the entire negative set (22382 nonobserved interactions with the lowest GO-based semantic measure proposed in this study).

**Figure 3.**
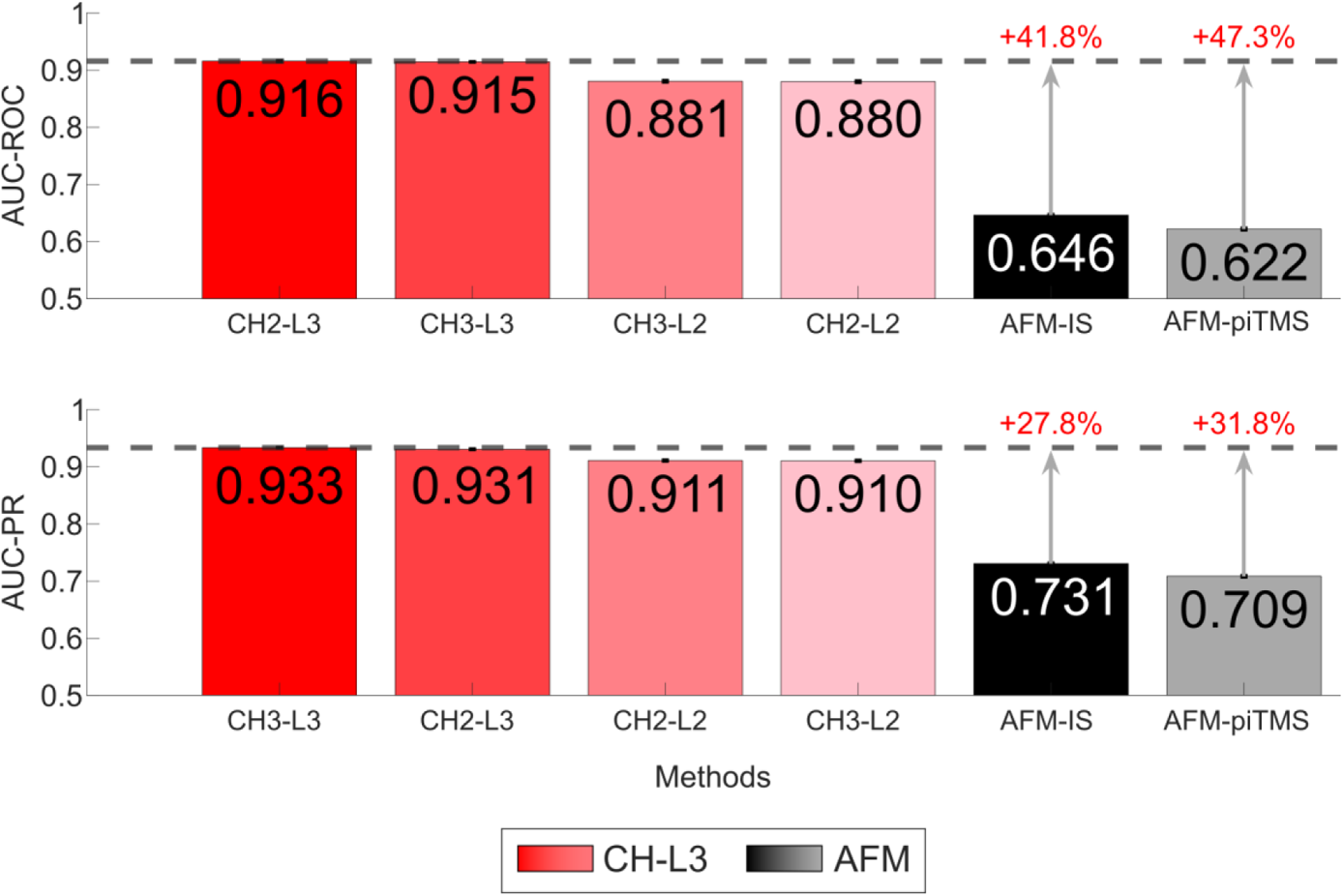
Comparing the performance of CH and AFM in predicting random 10%GSP+10%GSN protein interactions of the Yeast DIP network. Considering CH and AFM methods, the test involves predicting 10% positive and 10% negative interactions in the Yeast DIP network, which are randomly sampled from the GSP (Gold Standard Positives) and GSN (Gold Standard Negatives), respectively. To evaluate the performance, the links are sorted based on the predicted likelihood scores, and AUC-ROC and AUC-PR are computed for the entire sorted list of links. For each evaluation measure, the barplots report the mean performance (with error bars fwor standard error which is negligible) of each method on 10 realizations. The color of the CH bars varies depending on the performance value: the higher the performance is, the redder the CH bars appear. The arrows highlight the percentage of improvement between the best CH method and AFM methods as 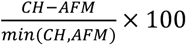.

The fourth important finding is that also in this BFEM evaluation framework the results are consistent with the ones provided in the previous section, indeed L3 methods performs better than L2 (Fig. 3, Suppl. Fig. 5-6). The fact that we provide BFEM as an effective GO-based evaluation framework is relevant because Kovács et al. reached contradictory results using a previously established GO-based framework^12,16^. Indeed, in Kovács et al., mere network topology prediction evaluation and GO evaluation disagree on what is the best strategy, with L2 methods performing better than L3^16^ in GO evaluation. Kovács et al. reported that the legitimacy of using GO terms or functional annotations to evaluate the quality of the predicted physical interactions should be questioned because they are based on similarity measures that are biased for L2. We show that BFEM does not suffer of this issue because, as demonstrated by Cannistraci (2018)^14^, we apply a more elaborated and stringent GO-filtered evaluation framework, which limits the analysis to the intersection of biological process (BP) and cellular component (CC) categories, as they provide more context and information about the protein interaction role within the cell. Indeed, proteins that do not participate to similar biological processes inside the same cellular compartments have less probability to interact. Furthermore, applying BFEM to a well-studied organism such as the yeast improves the quality and reliability of our results, to the point that we trust to investigate more in depth what are the reasons of the lower performance prediction of AFM.

We show that the set of interactions wrongly predicted by AFM is composed by proteins with amino acids that on average present significantly higher likelihood to belong to intrinsically disordered regions (IDR), and this is the fifth important finding. To this aim we introduce the interaction disorder index (IDI) which is a measure to quantify the presence of intrinsically disordered regions that affect the proteins involved in an interaction (see methods for details). Fig. 4a,c compare the distributions of IDI for true positive and false negative predictions in the entire GSP set (4476 interactions) considering AFM predictors, and report a significant difference (p-values are obtained applying a permutation test comparing the median of the data in the two groups). Fig. 4b,d report for AFM predictors a significant difference between the IDI distributions of true negative and false positive predictions in the entire GSN set (4476 interactions). This means that when AFM prediction is wrong most likely this is associated to the fact that the 3D structure of the proteins involved in the interactions is populated by IDR. The fact that AF has issues to predict the 3D structure of proteins with unstructured or intrinsically disordered regions is discussed in previous studies^1,49^, but our study is the first to propose a method to evaluate the extent to which this issue impacts also the prediction of protein interaction pairs. When we perform the same test for CH3-L3 (which is the best performing CH method), we notice (Fig.4 e,f) that all p-values are not significant and this implies that the prediction error (which is very low, see Fig. 3) of CH3-L3 is not associated with IDR. Finally, Fig.4 g,h report the remarkable computational time advantage of network shape intelligence (CH3-L3) over AFM prediction.

**Figure 4.**
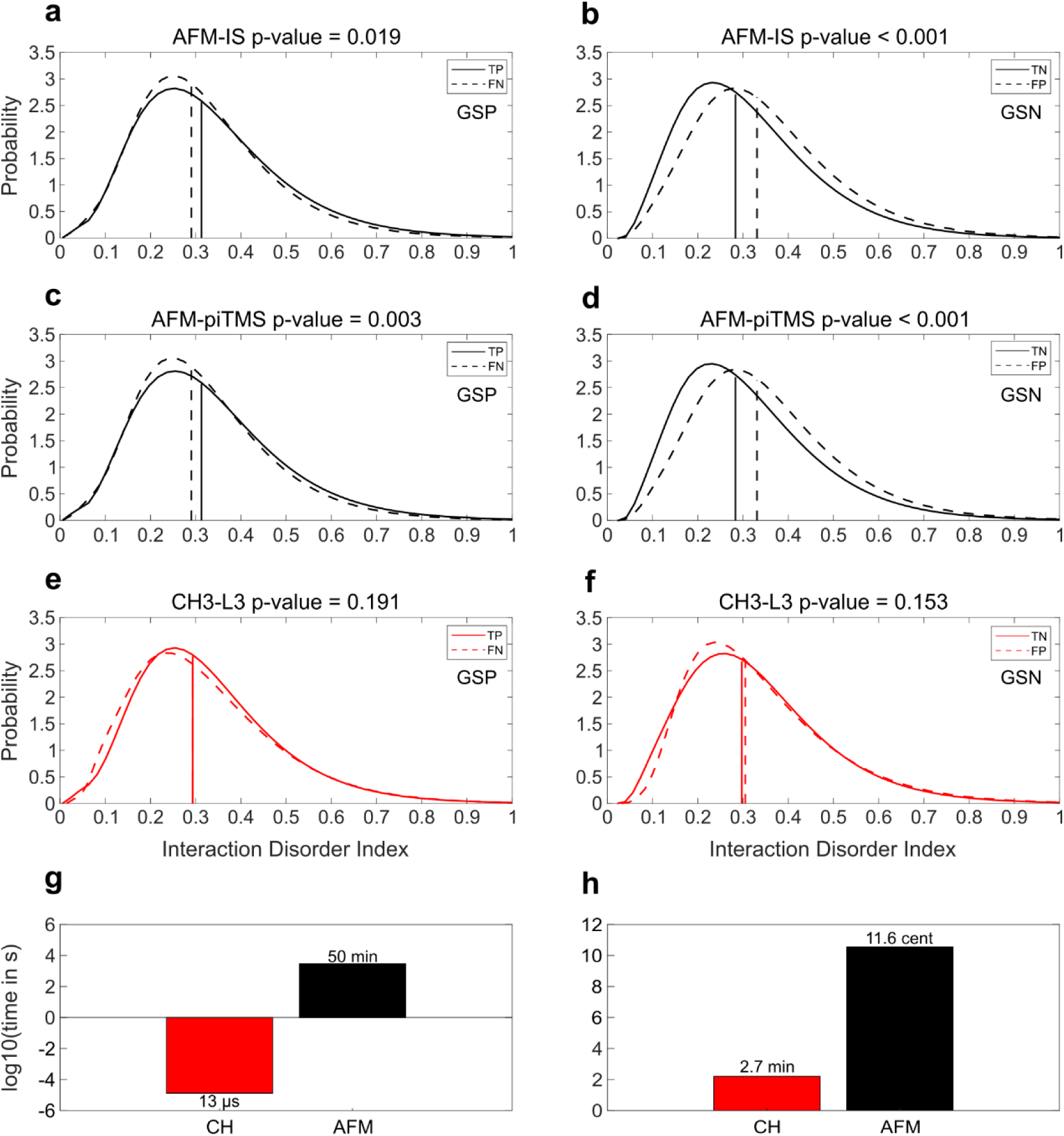
Interaction disorder index and computation time of CH and AFM. The probability density functions of the interaction disorder index for protein pairs in the entire (4476 interactions) golden standards positive (GSP, left panels) and the entire (4476 interactions) negative (GSN, right panels) sets predicted by AFM methods (**a**-**d**), and CH3-L3 (**e**-**f**). The solid line indicates correctly predicted interactions while the dashed line represents wrongly predicted interactions. The vertical line indicates the location of the median for each probability density function. (**g**) Average time to compute all variants of CH and AFM to predict one interaction. (**h**) Estimated time to compute CH variants and AFM variants to predict all the missing interactions. On top of each bar, min stands for minutes and cent for centuries.

## Discussion

Starting from our seminal studies on path-length 3 in bipartite networks^18^, passing through the L3 model by Kovács et al.^16^, in this study we show that L3 mathematical formula can emerge as part of a general theory for network automata modelling that through the lens of the Cannistraci-Hebb rule can produce results more astonishing than L3 itself for prediction of protein interactions. These results are confirmed in general complex network connectivity across different physical scales^24^.

The network automaton is a process that accesses to the local topology T(s) at state s and uses this topological memory trace (engram) to predict the next local connectivity T(s+1)^21^. Since the network represents a discrete memory of the state at time t of a complex dynamic systems whose rule of association between its generative latent variable is a manifold in a geometrical space^45,50–52^, the network automaton does not need training, because it reads directly from the network topology the knowledge that is necessary to decide how to evolve the connectivity at the next step.

Network shape intelligence methods such as Cannistraci-Hebb network automata for modelling and predicting network connectivity, using only local network information and without training, can outperform AlphaFold2 intelligence in protein interactions prediction, with a substantial advantage also in computational time. These results are gained by employing a bona fide evaluation methodology (BFEM) for assessment of protein interaction predictions, and the set of protein interactions mispredicted by AlphaFold2 contains amino acids that on average present significantly higher likelihood to belong to intrinsically disordered regions. To this aim we introduce the IDI which is a measure to quantify the presence of intrinsically disordered regions that affect the proteins involved in a single pairwise interaction however, as we comment in the method section, IDI can be extended to quantify the presence of intrinsically disordered regions in protein complexes. To advance modelling and prediction of protein interactions, next generation AlphaFold intelligence might integrate principles of network shape intelligence.

## Methods

### Dataset of protein-protein interactions

We evaluated our findings using 15 diverse interactomes spanning seven different species, as utilized by Kovács et al^16^. Additionally, we incorporated the Yeast DIP^29–31^ network to specifically compare the Network Shape Intelligence-based methods with the AlphaFold2 Intelligence approach. In Suppl. Tables 5-7, we report the main topological features and the sources with references to the associated studies of these networks.

The 15 interactomes include the following types. Systematic networks generated by a binary pipeline, such as the Human (HI-II-14), Yeast (Y2H-Union), and Arabidopsis (AI) interactomes. Literature-curated PPI networks of direct physical interactions, such as Lit-BM-13 and the BioGRID. From BioGRID interactome, we considered “direct interactions” and proteins assigned to the correct “taxid” corresponding to the studied species. Co-complex proteomic datasets, such as the Bioplex and Hein et al. AP-MS datasets. The Interactome3D dataset, which provides a summary of currently available interactions with structural evidence. For the Yeast DIP network, we obtained the list of PPIs (version Scere20170205) as pairs of Uniprot IDs and the corresponding FASTA sequences of the proteins (version DIP_20171201) from the Database of Interacting Proteins (DIP): https://dip.doe-mbi.ucla.edu/dip/Main.cgi. Once again, we specifically considered “direct interactions” and proteins assigned to the correct “taxid” corresponding to the yeast species.

Each PPI network was constructed and transformed into an undirected, unweighted format without self-loops, and only the largest connected component was retained.

### Construction of protein sets for the bona fide evaluation methodology (BFEM)

Applying a procedure adopted in previous studies^14,15^, the Wang Gene Ontology (GO) semantic similarity^53^ was computed for all protein pairs (existing and missing) in the Yeast DIP network, using the R package GOSemSim^54^. This method measures the similarity between GO terms and assigns a similarity score between 0 and 1 for each GO domain: biological process (BP), cellular component (CC) and molecular function (MF). Conversion from Uniprot IDs to Entrez IDs was performed with the Gene ID Conversion Tool of DAVID^55,56^, as the GOSemSim function requires a list of Entrez IDs as input.

To define the golden standard positive (GSP) set, we applied an elaborated and stringent GO-based framework as demonstrated by Cannistraci (2018)^14^, and limited our analysis to the biological process (BP) and cellular component (CC) categories as they provide more context and information about the protein’s role within the cell. This GO measure is based on the stringent criteria according to which two interacting proteins should share similar biological processes and be identified in similar cellular compartments^14^. Using Cannistraci’s GO similarity method, a GO similarity matrix with the minimum value between BP and CC for all existing protein interactions is created. Then, we sort these interactions by decreasing min(BP,CC) value, the higher the value the more at the top of the ranking. The rationale is that the most reliable interactions are the ones that rank at the top of the list even in the most conservative scenario in which the minimum value between BP and CC is considered. The positive set is composed by all the existing interactions (22382), and the golden standard positive (GSP) set is composed by the top 20% interactions (4476) in this ranking that were successfully predicted by AFM. To define the negative set, we introduce a new GO similarity measure according to which a GO similarity matrix with the maximum value between BP and CC for all missing interactions is created. Then, we sort these missing interactions by decreasing max(BP,CC) value, the lower the value the more at the bottom of the ranking. The rationale is that the less reliable interactions are the ones that rank at the bottom of the list even in the most relaxed scenario in which the maximum value between BP and CC is considered. The negative set is composed by selecting an amount of missing interactions at the bottom of this ranking that equals the number of existing interactions in the positive set (22382). The golden standard negative (GSN) set is composed by the bottom 20% interactions (4476) in this ranking that were successfully predicted by AFM. Please note that the final GO similarity matrix was smaller in size compared to the original adjacency matrix and its number of proteins and interactions is 4,672 and 21,816 respectively. This is due to reasons such as the failure to convert Uniprot IDs to Entrez IDs, or some proteins not having any annotations.

### AlphaFold2-Multimer (AFM) prediction of protein-protein interactions

In order to perform simulations in a reasonable amount of time, we used AFM-based ColabFold interface^57^ to model protein-protein interactions. Specifically, we utilized LocalColabFold (v1.3.0), an installer script that makes ColabFold functionality available on local machines, which speeds up single predictions by replacing AFM’s homology search with the faster MMseqs2: https://github.com/YoshitakaMo/localcolabfold. ColabFold was provided with multi-sequence FASTA files in input and executed with template mode on and number of recycles of 3 (default setting). The relaxation option was disabled as it was shown that AF2’s relaxation does not improve the accuracy of the predicted model^1^.

To predict heterodimeric protein complexes, we configured and tested six different settings using different combinations of the pair-mode (“unpaired+paired”, “unpaired”, “paired”) and the model-type (“AlphaFold2-multimer-v1”, “AlphaFold2-multimer-v2”). This resulted in six outputs for each prediction. We also modified ColabFold interface to include two metrics for estimating the confidence of a predicted complex as introduced by Gao et al.^6^: the predicted interface TM-score (AFM-piTMS) and the interface score (AFM-IS). Each predicted protein pair returned five (default) unrelaxed predicted structures in PDB format text file, a JSON format text file containing the times taken to run each section of the AFM pipeline, and another file collecting the AFM-based measures for each unrelaxed predicted structure.

The results of the comparison between the 6 settings are displayed in Suppl Fig. 7, and the setting that offered the best performance in the classification of the golden standard positive (GSP) and the golden standard negative (GSN) protein pairs of the Yeast DIP network is ‘unpaired+paired’, therefore it was used to compare the performance with the Network Shape Intelligence methods in the Fig. 3,4 and Suppl. Fig. 5,6 of this study. In this study, for a fair comparison with the network shape intelligence methods, we decided to adopt Gao et al.^6^ method because, unlike Evans’ piTM score^3^ (named ipTM score in their paper), Gao’s piTM score has an adjusted normalization factor in the formula to better deal with the cases where a low number of contacts at the interface are observed. Besides, Gao et al. interface score (IS) is similar to Evans’ ipTM score, however the IS has the advantage to be calculated for each protein chain separately by considering the observed number of interface residues in that chain. The optimal local reference for calculating the score for a specific chain is selected using interface residues that do not belong to that chain. Gao et al. approach allows for a more targeted analysis of interface residues (versus full chain in Evans et al.), which is particularly relevant for protein interaction predictions, which is the focus of this study. Besides, we preferred to adopt Gao et al.^6^ method instead of Bryant et al.^5^ method (predicted DockQ score: pDockQ) because Gao et al. scoring methods (piTM score and interface score IS) are specifically designed to evaluate protein-protein interactions and provide a comprehensive analysis of interface residues. On the other hand, Bryant et al. method (pDockQ) focuses on estimating the overall quality of the predicted model rather than quantifying the quality of each individual interface within a multi-chain complex. One advantage of Gao et al.’s piTM score over pDockQ is that it incorporates an adjusted normalization factor in its formula. As we explained above, this factor accounts for situations where a low number of contacts at the interface are observed. By considering this normalization factor, Gao et al.’s piTM score provides a more accurate assessment of the interface quality, particularly in cases where the number of interface residues is limited. In contrast, pDockQ does not have such a normalization factor, which may lead to biased evaluations in scenarios with few interface contacts. Moreover, the IS of Gao et al. is similar to Bryant et al. pDockQ in that they both evaluate the quality of the interface. However, IS offers the advantage of calculating the score for each protein chain separately. This individual assessment of each chain’s interface allows for a more targeted analysis of the interface residues, taking into account the observed number of residues specific to each chain. Gao et al. achieve this by selecting the optimal local reference using interface residues that do not belong to the chain being evaluated. In contrast, Bryant et al. pDockQ does not provide a chain-specific analysis and evaluates the overall interface quality without considering the contributions of individual chains. Therefore, based on the adjusted normalization factor incorporated in Gao et al. piTM score and the chain-specific evaluation provided by Gao et al. IS, we have opted to employ Gao et al. scoring methods over Bryant et al. pDockQ for our research work, as they align more closely with our specific goals and requirements in the analysis of protein-protein interactions within multi-chain complexes.

### Cannistraci-Hebb network automata

The Cannistraci-Hebb (CH) theory^21,24,26,28^ has been introduced as a revision of the local-community paradigm (LCP) theory^14,15,18,22^ and it has been formalized within the framework of network automata^21,24,26,28^. While the LCP paradigm emphasized the importance to complement the information related to the common neighbours with the interactions between them (internal local-community-links), the CH rule is based on the local isolation of the common neighbours by minimizing their interactions external to the local community (external local-community links). CH network automata on paths of length n are all the network automata models that explicitly consider the minimization of the external local community-links within a local community characterized by paths of length n^26^. We considered the following CH models: CH1-L2/L3, CH2-L2/L3, CH3-L2/L3; also RA-L2/L3 is considered a CH model because its definition relies on the minimization of external links in the local community. In addition, each of the four CH models applied the associated CH-SPcorr score for subranking^26^, in order to internally sub-rank all the node pairs characterized by the same CH score, reducing the ranking uncertainty of node pairs that are tied-ranked. The code to run these methods is available at the web address provided in the code availability section of the study.

### Interaction disorder index (IDI)

The interaction disorder index (IDI) is a measure that we introduce to quantify the extent to which intrinsically disordered regions occur and affect the proteins involved in an interaction. The procedure to compute the IDI is the following. We employ the IUPred3 algorithm^58^: a widely used computational tool that predicts the likelihood of each amino acid in a protein sequence being in a disordered region. IUPred3 is applied to the amino acids of each protein within an interaction pair, and the mean of amino acid IUPred3 scores for protein 1 and protein 2 are computed, for instance S_1_ is the mean of the scores for protein 1 and S_2_ for protein 2 respectively. To calculate the value of interaction disorder index for a protein interaction, we take the average of S_1_ and S_2_: 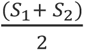. To calculate the value of interaction disorder index for a protein complex, or for an ensemble of interactions in a network, we suggest adopting a centrality measure (for instance, the mean or median) of the probability distribution formed by the IDIs values of the interactions in the complex or in the interaction ensemble.

In Figure 4, the p-values for each panel are obtained applying a permutation test comparing the median (we prefer the median to the mean because the distributions are skewed) of the data in the two groups. The probability density functions are plotted using a kernel smoothing function (ksdensity in MATLAB) with a bandwidth value of 0.3 because this bandwidth value offers a distributions shape that reflects the statistics of the data obtained by the permutation test.

### Time computation of CHA and AFM on the Yeast DIP network

To determine the average computation time of CH for predicting one interaction, we annotated the time required for CH to predict all the node pairs and divided by the total number of pairs. For AFM we annotated the time to predict a random sample of 1% PPIs and divided that by the total number of predicted interactions. To determine the computational time of CH or AFM for predicting all missing interactions, we multiplied the average prediction time of one interaction for all the missing interactions. Note that the CH algorithm was executed in parallel using 12 workers on a local workstation (Hardware and software section), while AFM simulations were launched on the multi-GPU cluster at the Center for Information Services and High-Performance Computing (ZIH) of TU Dresden (see below Hardware and software section).

### Hardware and software

The CH network automata computation were executed in a workstation under Windows 10 Enterprise 2016 LTSB with 176 GB RAM, and 2 Intel(R) Xeon(R) CPU X5550 processors (4 cores each) with 2.67 GHz. The software environment is MATLAB 2019a.

The AFM simulations were executed on the multi-GPU cluster “Alpha Centauri” at the ZIH of the TU Dresden. It has 34 nodes, each with:

- 8 x NVIDIA A100-SXM4 (40 GB RAM)
- 2 x AMD EPYC 7352 CPU (24 cores) with 2.3 GHz with multi-threading enabled
- 1 TB RAM

One GPU and eight CPU cores were used to run AFM properly. The loaded modules environment in the cluster are GCC/10.2.0, CUDA/11.3.1, OpenMPI/4.0.5. The software environment is Python 3.9.

### Data availability

The networks to reproduce the result of this study are provide at the GitHub repository: https://github.com/biomedical-cybernetics/NSIforPPI [they will be released after article acceptance].

### Code Availability

The MATLAB codes to compute CH network automata and to reproduce the results of some main figures of the study are publicly available at the GitHub repository: https://github.com/biomedical-cybernetics/NSIforPPI [they will be released after article acceptance].

## Acknowledgements

We thank: Yi Huang (Tsinghua University) and Daniele Macuglia (Peking University) for the comments and for proofreading the article; the Center for Information Services and High Performance Computing (ZIH) of the TU Dresden and ScaDS.AI for providing HPC resources; the BIOTEC System Administrators for their IT support; Sabine Zeissig for the administrative assistance at BIOTEC; Yue Wu, YuanYuan Song, Yining Xin, Giada Zhou, Shao Qian Yi, Lixia Huang, and Weijie Guan for the administrative support at THBI; Hao Pang for the IT support at THBI.

## Author Contributions

CVC proposed the definition of Network Shape Intelligence (NSI) and conceived the theory, the experiments and the content of the study. AM and CVC designed the algorithm, the computational experiments, figures and items for the CH network automata section, and IA realized them. IA and CVC designed the algorithm, the computational experiments, figures and items for the CH network automata versus AFM comparison section, and IA realized them. CVC invented the BFEM. IA hypothesized the relation between IDR and mis-predictions of AFM, and CVC designed the statistical procedure to prove it. IA performed the computational experiments and prepared the code to implement them. ML and UM optimized the setup of AlfaFold2 on high performance computing systems and DMR guided IA in job submission. IA and CVC analyzed and interpreted the results. CVC wrote the article and the supplementary information including notes of IA, and corrections from the other authors. CVC planned, directed and supervised the study.

