## Supplementary Information for "Network shape intelligence outperforms AlphaFold2 intelligence in vanilla protein interaction prediction"

† First co-authorship

### Supplementary Information

### Supplementary Note 1. CH network automata generalization in path n and sub-ranking strategy

#### *CH network automata generalization on path of length n (ln).*

To formulate CH1, CH2 and CH3 in paths of length *n (ln)*, we need to define what the local community is in paths of length *n*. To this aim, it is easier to provide an example for paths of length three (*l3*) as represented in the Fig. 1. The common neighbours are all the intermediate nodes touched by any path of length three that connects the two seed nodes *u* and *v*. The iLCL are the links between common neighbours, to guarantee a direct flux of information that forms a local-tunnel between the seed nodes (Fig 1). The local community is the structure that is posed at the centre of the tunnel formed by the L3 paths and it is composed of the common neighbours and the iLCL. Hence, the mathematical formulae for the CH1, CH2 and CH3 models for paths of length *n* are:

$$CH1\_Ln(u,v) = \sum_{z_1 \dots z_{n-1} \in ln(u,v)} \frac{\left(di_{z_1} * \dots * di_{z_{n-1}}\right)^{\frac{1}{n-1}}}{\left(d_{z_1} * \dots * d_{z_{n-1}}\right)^{\frac{1}{n-1}}}$$

$$CH2\_Ln(u,v) = \sum_{z_1 \dots z_{n-1} \in \ln(u,v)} \frac{\left(di^*_{z_1} * \dots * di^*_{z_{n-1}}\right)^{\frac{1}{n-1}}}{\left(de^*_{z_1} * \dots * de^*_{z_{n-1}}\right)^{\frac{1}{n-1}}}$$

$$CH3\_Ln(u,v) = \sum_{z_1 \dots z_{n-1} \in \ln(u,v)} \frac{1}{\left(de^*_{z_1} * \dots * de^*_{z_{n-1}}\right)^{\frac{1}{n-1}}}$$

where: $u$ and $v$ are the two seed nodes of the candidate interaction; $z_1 \dots z_{n-1}$ are the intermediate nodes (common neighbours) on the considered path of length $n$ ($ln$); $d_{z_1} \dots d_{z_{n-1}}$ are the respective node degrees; $di_{z_1} \dots di_{z_{n-1}}$ are the respective internal node degrees (number of iLCL); $de_{z_1} \dots de_{z_{n-1}}$ are the respective external node degrees (number of eLCL); terms with an asterisk as superscript indicate that a unitary value is added ($di^*_{z_1} = 1 + di_{z_1}$ and $de^*_{z_1} = 1 + de_{z_1}$); and the summation is executed over all the paths of length $n$. All the formulae for L2, L3 and Ln are summarized in Fig. 1 and Suppl. Table 1.

We stress that till now we never spoke about the triadic closure principle[1] because we believe that common neighbours are not associated to any triadic closure in general, and this might be a misleading principle. In our opinion, a correct principle to define common neighbours is in relation to the definition of local paths of length $n$. The common neighbours are in general, according to our proposed definition, all the intermediate nodes touched by any path of length $n$ between two nodes in the network. In case of paths of length 2 ($l2$) this is specifically coincident with triadic closure, but in case of paths of length $n$ we propose to adopt the general definition: 'local ring' closure[2].

*Sub-ranking strategy*

First, it adjusts a bias in performance evaluation between L2 and L3 predictors because L2 predictors assign likelihood zero to all nonadjacent node pairs that do not have a mere triadic closure, whereas L3 predictors are including also quadratic closure. Second, and more in general, it adjusts a bias in performance evaluation between local topology methods (such as the automata considered in this study) that assigns a likelihood zero to any nonadjacent node pairs that are not involved in a topological neighbourhood, and global topology methods that assign a likelihood to all nonadjacent node pair. The details on the sub-ranking strategy are provided in Muscoloni et al.[3]

| **Epitopological rationale** | **L2** | **L3** | **Ln** |
|---|---|---|---|
| **Resource Allocation (RA)**<br>*degree penalty* | $\frac{1}{d_z}$ | $\frac{1}{\sqrt{d_{z_1} * d_{z_2}}}$ | $\frac{1}{\left(d_{z_1} * \dots * d_{z_{n-1}}\right)^{\frac{1}{n-1}}}$ |
| **Cannistraci-Hebb 1 (CH1)**<br>*Internal degree reward and external degree penalty* | $\frac{di_z}{d_z}$ | $\frac{\sqrt{di_{z_1} * di_{z_2}}}{\sqrt{d_{z_1} * d_{z_2}}}$ | $\frac{\left(di_{z_1} * \dots * di_{z_{n-1}}\right)^{\frac{1}{n-1}}}{\left(d_{z_1} * \dots * d_{z_{n-1}}\right)^{\frac{1}{n-1}}}$ |

| | | | |
|---|---|---|---|
| **Cannistraci-Hebb 2 (CH2)**<br>*Internal degree reward and external degree penalty* | $\frac{di_z^*}{de_z^*} = \frac{1+di_z}{1+de_z}$ | $\frac{\sqrt{di_{z_1}^* * di_{z_2}^*}}{\sqrt{de_{z_1}^* * de_{z_2}^*}}$ | $\frac{(di_{z_1}^* * \dots * di_{z_{n-1}}^*)^{\frac{1}{n-1}}}{(de_{z_1}^* * \dots * de_{z_{n-1}}^*)^{\frac{1}{n-1}}}$ |
| **Cannistraci-Hebb 3 (CH3)**<br>*external degree penalty* | $\frac{1}{de_z^*} = \frac{1}{1+de_z}$ | $\frac{1}{\sqrt{de_{z_1}^* * de_{z_2}^*}}$ | $\frac{1}{(de_{z_1}^* * \dots * de_{z_{n-1}}^*)^{\frac{1}{n-1}}}$ |
| **iLCL**<br>*internal degree reward* | $\frac{di_z}{di_z^*} = \frac{di_z}{1+di_z}$ | $\frac{\sqrt{di_{z_1} * di_{z_2}}}{\sqrt{di_{z_1}^* * di_{z_2}^*}}$ | $\frac{(di_{z_1} * \dots * di_{z_{n-1}})^{\frac{1}{n-1}}}{(di_{z_1}^* * \dots * di_{z_{n-1}}^*)^{\frac{1}{n-1}}}$ |

| **Network automata mechanistic model** | | |
|---|---|---|
| **L2** | **L3** | **Ln** |
| $RA\_L2(u,v) = \sum_{z \in l2} \frac{1}{d_z}$ | $RA\_L3(u,v) = \sum_{z_1,z_2 \in l3} \frac{1}{\sqrt{d_{z_1} * d_{z_2}}}$ | $RA\_Ln(u,v) = \sum_{z_1 \dots z_{n-1} \in ln} \frac{1}{(d_{z_1} * \dots * d_{z_{n-1}})^{\frac{1}{n-1}}}$ |
| $CH1\_L2(u,v) = \sum_{z \in l2} \frac{di_z}{d_z}$ | $CH1\_L3(u,v) = \sum_{z_1,z_2 \in l3} \frac{\sqrt{di_{z_1} * di_{z_2}}}{\sqrt{d_{z_1} * d_{z_2}}}$ | $CH1\_Ln(u,v) = \sum_{z_1 \dots z_{n-1} \in ln} \frac{(di_{z_1} * \dots * di_{z_{n-1}})^{\frac{1}{n-1}}}{(d_{z_1} * \dots * d_{z_{n-1}})^{\frac{1}{n-1}}}$ |
| $CH2\_L2(u,v) = \sum_{z \in l2} \frac{di_z^*}{de_z^*}$ | $CH2\_L3(u,v) = \sum_{z_1,z_2 \in l3} \frac{\sqrt{di_{z_1}^* * di_{z_2}^*}}{\sqrt{de_{z_1}^* * de_{z_2}^*}}$ | $CH2\_Ln(u,v) = \sum_{z_1 \dots z_{n-1} \in ln} \frac{(di_{z_1}^* * \dots * di_{z_{n-1}}^*)^{\frac{1}{n-1}}}{(de_{z_1}^* * \dots * de_{z_{n-1}}^*)^{\frac{1}{n-1}}}$ |
| $CH3\_L2(u,v) = \sum_{z \in l2} \frac{1}{de_z^*}$ | $CH3\_L3(u,v) = \sum_{z_1,z_2 \in l3} \frac{1}{\sqrt{de_{z_1}^* * de_{z_2}^*}}$ | $CH3\_Ln(u,v) = \sum_{z_1 \dots z_{n-1} \in ln} \frac{1}{(de_{z_1}^* * \dots * de_{z_{n-1}}^*)^{\frac{1}{n-1}}}$ |
| $iLCL\_L2(u,v) = \sum_{z \in l2} \frac{di_z}{di_z^*}$ | $iLCL\_L3(u,v) = \sum_{z_1,z_2 \in l3} \frac{\sqrt{di_{z_1} * di_{z_2}}}{\sqrt{di_{z_1}^* * di_{z_2}^*}}$ | $iLCL\_Ln(u,v) = \sum_{z_1 \dots z_{n-1} \in ln} \frac{(di_{z_1} * \dots * di_{z_{n-1}})^{\frac{1}{n-1}}}{(di_{z_1}^* * \dots * di_{z_{n-1}}^*)^{\frac{1}{n-1}}}$ |

**Suppl. Table 1. Mathematical description of the methods.**

Notation: $u, v$ are the seed nodes; $z$ is the intermediate node; $d_z$ is the degree of $z$; $di_z$ is the internal degree (number of iLCL) of $z$; $de_z$ is the external degree (number of eLCL) of $z$. For any degree it is valid the following: $d^* = 1 + d$.

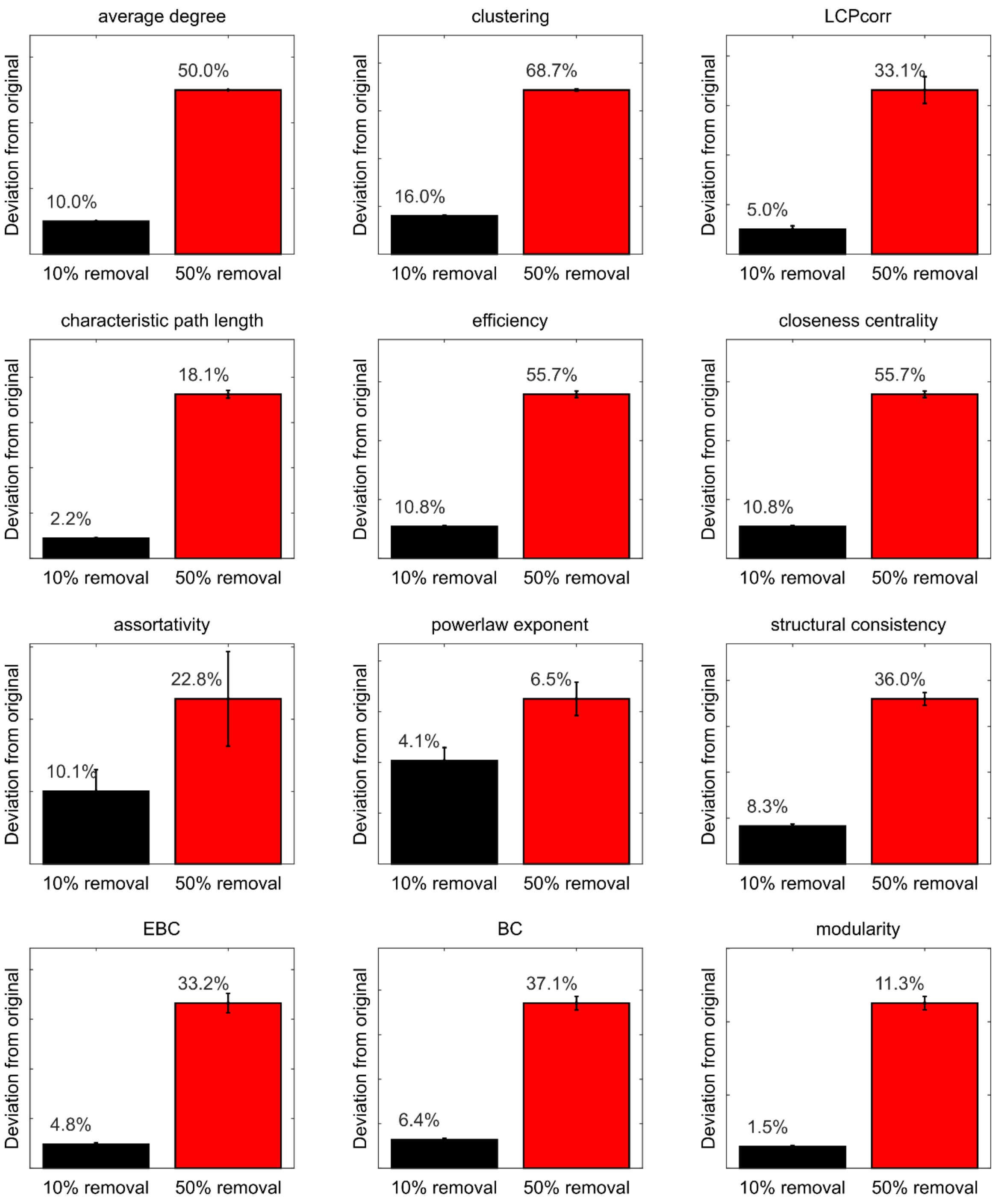


**Suppl. Fig. 1. Relative deviation of topological measures for PPI networks.**

For each PPI network, 10 reduced networks have been obtained randomly removing 10% of the links and 10 networks randomly removing 50% of the links. For the original network and for each of the reduced ones, several topological measures have been computed. The barplots report for each topological measure the mean relative deviation of the measure computed in the reduced network with respect to the original network, comparing the 10% and 50% link removal. The relative deviation is computed as $|(x_{original} - x_{reduced})/x_{original}|$ and the reported value is the average over all the PPI networks and 10 repetitions.

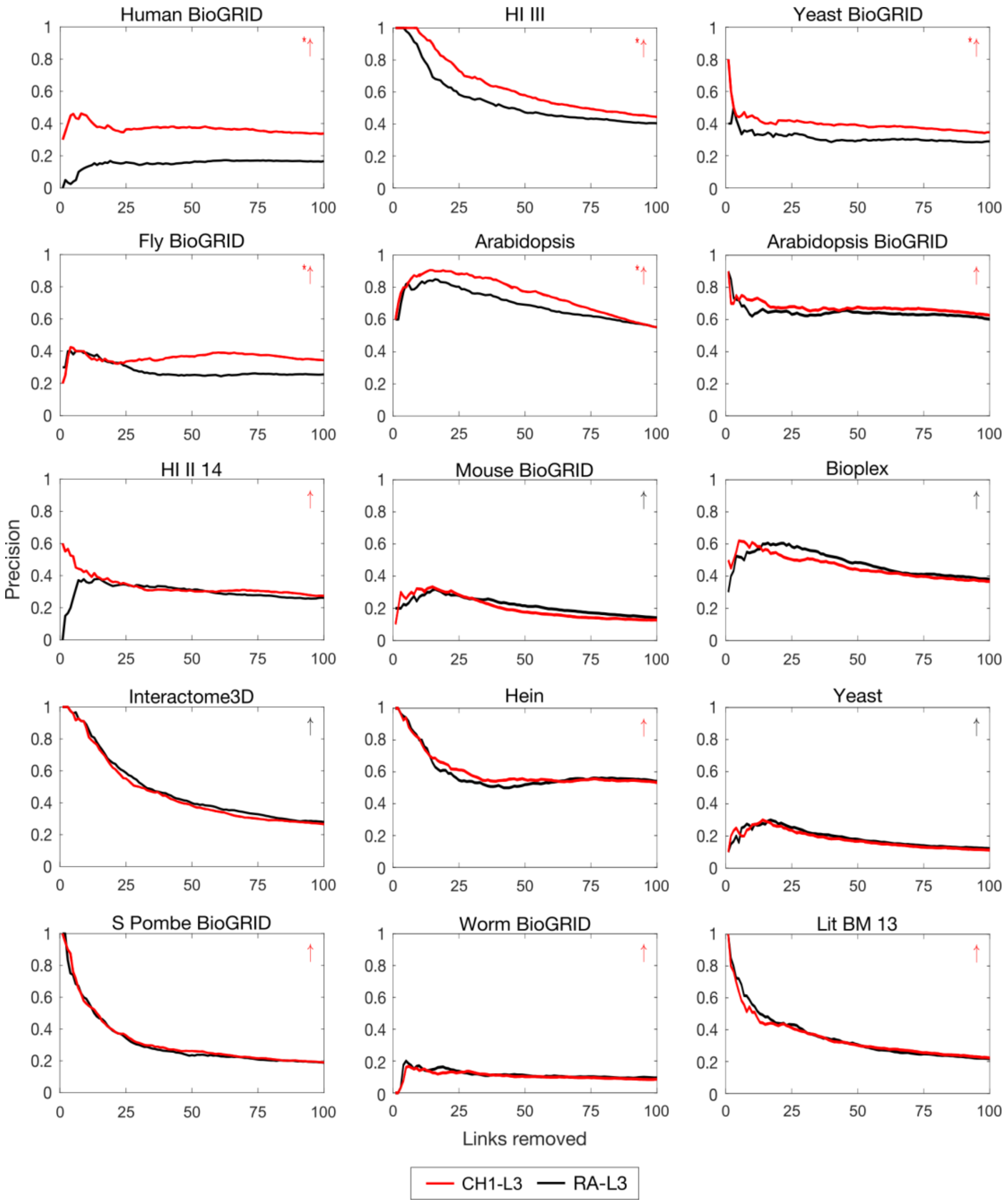


**Suppl. Fig. 2. Precision curve evaluation for PPI networks: CH1-L3 vs RA-L3 methods.**

For each PPI network, 10% of links were randomly removed (10 repetitions) and the algorithms were executed to assign likelihood scores to the nonobserved links in these reduced networks. To evaluate the performance, the links are ranked by likelihood scores and the precision is computed as the percentage of removed links among the top-r in the ranking, for each r from 1 up to 100 at steps of 1. The plots report for each network the precision curve (averaged over the 10 repetitions) for the L3-based link prediction methods RA-L3 and CH1-L3. For each network, a red or black arrow is reported on the top-right of the subplot to indicate respectively if the average area under precision curve (AUP@100) is higher or lower for CH1-L3 with respect to RA-L3. A permutation test for the mean AUP@100 has been computed, and an asterisk is reported next to the arrow in case of statistical difference (p-value ≤ 0.05). Error bars are not shown because negligible. The networks are ordered (top-down, left-right) by decreasing absolute difference of mean AUP@100 and, in case of tie, by increasing p-value between the two methods.

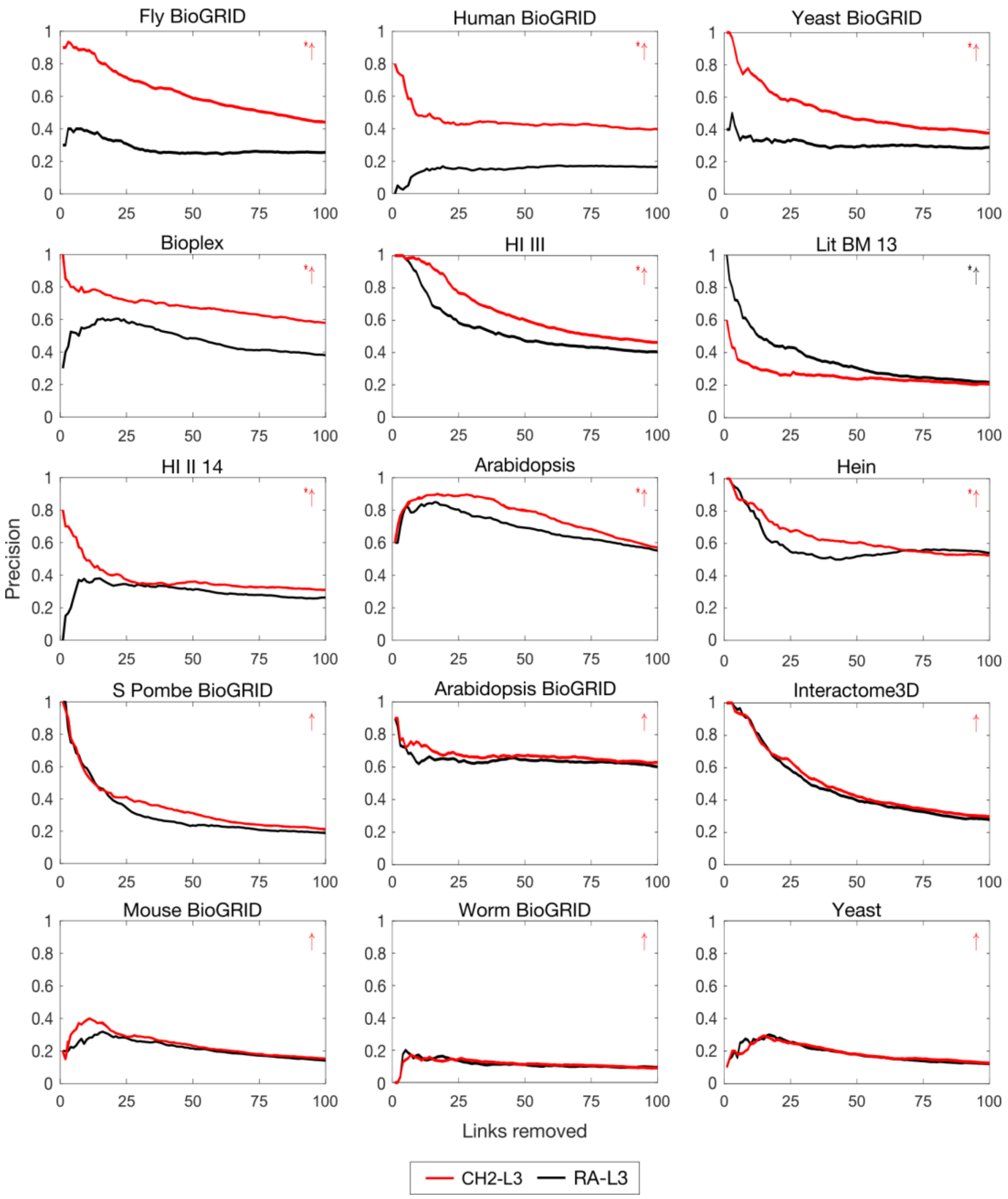


**Suppl. Fig. 3. Precision curve evaluation for PPI networks: CH2-L3 vs RA-L3 methods.**

For each PPI network, 10% of links were randomly removed (10 repetitions) and the algorithms were executed to assign likelihood scores to the non-observed links in these reduced networks. In order to evaluate the performance, the links are ranked by likelihood scores and the precision is computed as the percentage of removed links among the top-r in the ranking, for each r from 1 up to 100 at steps of 1. The plots report for each network the precision curve (averaged over the 10 repetitions) for the L3-based link prediction methods RA-L3 and CH2-L3. For each network, a red or black arrow is reported on the top-right of the subplot to indicate respectively if the average area under precision curve (AUP@100) is higher or lower for CH2-L3 with respect to RA-L3. A permutation test for the mean AUP@100 has been computed, and an asterisk is reported next to the arrow in case of statistical difference (p-value $\leq 0.05$). Error bars are not shown because negligible. The networks are ordered (top-down, left-right) by decreasing absolute difference of mean AUP@100 and, in case of tie, by increasing p-value between the two methods.

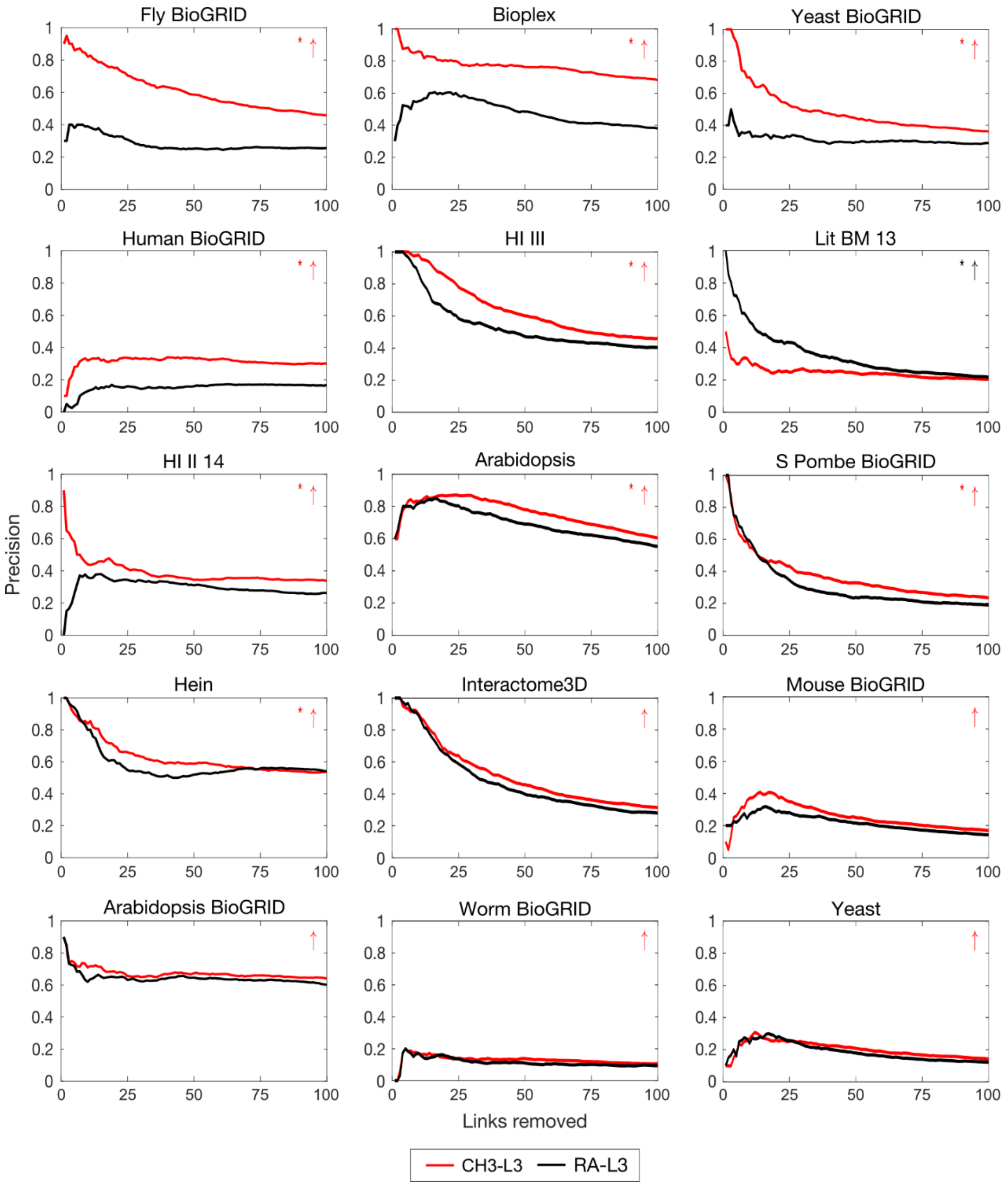


**Suppl. Fig. 4. Precision curve evaluation for PPI networks: CH3-L3 vs RA-L3 methods.**

For each PPI network, 10% of links were randomly removed (10 repetitions) and the algorithms were executed to assign likelihood scores to the non-observed links in these reduced networks. In order to evaluate the performance, the links are ranked by likelihood scores and the precision is computed as the percentage of removed links among the top-r in the ranking, for each r from 1 up to 100 at steps of 1. The plots report for each network the precision curve (averaged over the 10 repetitions) for the L3-based link prediction methods RA-L3 and CH3-L3. For each network, a red or black arrow is reported on the top-right of the subplot to indicate respectively if the average area under precision curve (AUP@100) is higher or lower for CH3-L3 with respect to RA-L3. A permutation test for the mean AUP@100 has been computed, and an asterisk is reported next to the arrow in case of statistical difference (p-value ≤ 0.05). Error bars are not shown because negligible. The networks are ordered (top-down, left-right) by decreasing absolute difference of mean AUP@100 and, in case of tie, by increasing p-value between the two methods.

| | CH3 L3 | CH2 L3 | CH1 L3 | RA L3 | iLCL L3 | CH1 L2 | CH2 L2 | iLCL L2 | CH3 L2 | RA L2 |
|---|---|---|---|---|---|---|---|---|---|---|
| **mean ranking** | 1.80 | 1.90 | 3.20 | 4.20 | 4.60 | 6.00 | 6.60 | 8.07 | 8.80 | 9.83 |
| **mean AUP** | 0.47 | 0.47 | 0.41 | 0.38 | 0.37 | 0.22 | 0.21 | 0.15 | 0.10 | 0.06 |
| *S_Pombe_BioGRID* | 0.37 | 0.35 | 0.32 | 0.31 | 0.30 | 0.16 | 0.12 | 0.05 | 0.06 | 0.03 |
| *Yeast* | 0.20 | 0.19 | 0.18 | 0.19 | 0.16 | 0.08 | 0.05 | 0.06 | 0.03 | 0.02 |
| *Mouse_BioGRID* | 0.26 | 0.24 | 0.20 | 0.22 | 0.16 | 0.05 | 0.05 | 0.04 | 0.04 | 0.02 |
| *Worm_BioGRID* | 0.13 | 0.12 | 0.11 | 0.11 | 0.10 | 0.08 | 0.06 | 0.06 | 0.04 | 0.04 |
| *HI_II_14* | 0.39 | 0.38 | 0.33 | 0.30 | 0.31 | 0.24 | 0.19 | 0.20 | 0.07 | 0.05 |
| *Arabidopsis* | 0.76 | 0.77 | 0.76 | 0.70 | 0.75 | 0.31 | 0.21 | 0.18 | 0.05 | 0.02 |
| *Yeast_BioGRID* | 0.49 | 0.52 | 0.39 | 0.31 | 0.28 | 0.18 | 0.21 | 0.11 | 0.04 | 0.02 |
| *Interactome3D* | 0.52 | 0.50 | 0.46 | 0.48 | 0.45 | 0.17 | 0.16 | 0.09 | 0.08 | 0.03 |
| *Lit_BM_13* | 0.25 | 0.26 | 0.35 | 0.35 | 0.30 | 0.11 | 0.10 | 0.08 | 0.06 | 0.05 |
| *Hein* | 0.63 | 0.63 | 0.60 | 0.59 | 0.59 | 0.47 | 0.51 | 0.43 | 0.44 | 0.33 |
| *Fly_BioGRID* | 0.62 | 0.62 | 0.36 | 0.28 | 0.27 | 0.07 | 0.03 | 0.06 | 0.01 | 0.01 |
| *Arabidopsis_BioGRID* | 0.67 | 0.67 | 0.67 | 0.64 | 0.67 | 0.15 | 0.19 | 0.00 | 0.02 | 0.02 |
| *HI_III* | 0.65 | 0.66 | 0.63 | 0.54 | 0.58 | 0.42 | 0.42 | 0.34 | 0.13 | 0.10 |
| *Bioplex* | 0.77 | 0.68 | 0.46 | 0.48 | 0.43 | 0.62 | 0.68 | 0.32 | 0.42 | 0.15 |
| *Human_BioGRID* | 0.31 | 0.45 | 0.37 | 0.15 | 0.22 | 0.26 | 0.20 | 0.19 | 0.07 | 0.06 |

**Suppl. Table 2. AUP evaluation of top-100 (AUP@100) interactions for PPI networks.**

For each network 10% of links have been randomly removed (10 repetitions) and the algorithms have been executed in order to assign likelihood scores to the non-observed links in these reduced networks. In order to evaluate the performance, the links are ranked by likelihood scores and the area under precision curve (AUP@100) is computed for the top-100 links in the ranking. The table reports for each network the mean AUP@100 over the random repetitions. The first rows show the mean precision and the mean ranking over the entire dataset. The networks are sorted by increasing number of nodes *N*.

| | CH3 L3 | CH2 L3 | CH1 L3 | RA L3 | iLCL L3 | CH2 L2 | CH1 L2 | CH3 L2 | iLCL L2 | RA L2 |
|---|---|---|---|---|---|---|---|---|---|---|
| **mean ranking** | 1.97 | 2.03 | 3.13 | 3.43 | 4.77 | 6.13 | 6.80 | 8.63 | 8.80 | 9.30 |
| **mean AUC-mROC** | 0.76 | 0.76 | 0.75 | 0.75 | 0.74 | 0.69 | 0.68 | 0.66 | 0.66 | 0.65 |
| *S_Pombe_BioGRID* | 0.76 | 0.76 | 0.76 | 0.76 | 0.75 | 0.67 | 0.66 | 0.64 | 0.62 | 0.63 |
| *Yeast* | 0.70 | 0.70 | 0.70 | 0.70 | 0.69 | 0.62 | 0.61 | 0.61 | 0.60 | 0.61 |
| *Mouse_BioGRID* | 0.72 | 0.72 | 0.71 | 0.71 | 0.69 | 0.63 | 0.62 | 0.62 | 0.61 | 0.62 |
| *Worm_BioGRID* | 0.69 | 0.68 | 0.68 | 0.68 | 0.67 | 0.62 | 0.61 | 0.61 | 0.60 | 0.61 |
| *HI_II_14* | 0.75 | 0.75 | 0.74 | 0.73 | 0.73 | 0.69 | 0.69 | 0.64 | 0.68 | 0.64 |
| *Arabidopsis* | 0.81 | 0.81 | 0.81 | 0.80 | 0.80 | 0.71 | 0.72 | 0.67 | 0.69 | 0.66 |
| *Yeast_BioGRID* | 0.76 | 0.76 | 0.74 | 0.73 | 0.72 | 0.71 | 0.70 | 0.66 | 0.68 | 0.65 |
| *Interactome3D* | 0.80 | 0.80 | 0.79 | 0.80 | 0.78 | 0.73 | 0.72 | 0.70 | 0.69 | 0.68 |
| *Lit_BM_13* | 0.73 | 0.73 | 0.74 | 0.75 | 0.73 | 0.68 | 0.67 | 0.66 | 0.66 | 0.66 |
| *Hein* | 0.78 | 0.78 | 0.78 | 0.78 | 0.77 | 0.75 | 0.75 | 0.73 | 0.73 | 0.72 |
| *Fly_BioGRID* | 0.76 | 0.76 | 0.72 | 0.72 | 0.72 | 0.60 | 0.62 | 0.59 | 0.61 | 0.59 |
| *Arabidopsis_BioGRID* | 0.80 | 0.80 | 0.79 | 0.79 | 0.79 | 0.71 | 0.70 | 0.66 | 0.65 | 0.65 |
| *HI_III* | 0.78 | 0.78 | 0.77 | 0.77 | 0.77 | 0.72 | 0.72 | 0.67 | 0.71 | 0.67 |
| *Bioplex* | 0.79 | 0.78 | 0.75 | 0.75 | 0.74 | 0.76 | 0.75 | 0.73 | 0.71 | 0.69 |
| *Human_BioGRID* | 0.73 | 0.75 | 0.74 | 0.71 | 0.71 | 0.70 | 0.70 | 0.66 | 0.69 | 0.66 |

**Suppl. Table 3. AUC-mROC evaluation of all non-observed interactions for PPI networks.**
For each network 10% of links have been randomly removed (10 repetitions) and the algorithms have been executed in order to assign likelihood scores to the non-observed links in these reduced networks. In order to evaluate the performance, the links are ranked by likelihood scores and the AUC-mROC is computed for all the non-observed links in the ranking. The table reports for each network the mean AUC-mROC over the random repetitions. The first rows show the mean AUC-mROC and the mean ranking over the entire dataset. The networks are sorted by increasing number of nodes *N*.

| | CH3 L3 | CH2 L3 | RA L3 | CH1 L3 | iLCL L3 | CH2 L2 | CH1 L2 | iLCL L2 | CH3 L2 | RA L2 |
|---|---|---|---|---|---|---|---|---|---|---|
| **mean ranking** | 1.13 | 2.20 | 2.90 | 3.77 | 5 | 6.53 | 6.67 | 8.63 | 8.83 | 9.33 |
| **mean AUC-PR** | 0.050 | 0.044 | 0.042 | 0.039 | 0.032 | 0.011 | 0.011 | 0.007 | 0.006 | 0.005 |
| *S_Pombe_BioGRID* | 0.054 | 0.049 | 0.044 | 0.042 | 0.034 | 0.008 | 0.008 | 0.002 | 0.005 | 0.004 |
| *Yeast* | 0.021 | 0.019 | 0.019 | 0.017 | 0.014 | 0.003 | 0.003 | 0.002 | 0.002 | 0.002 |
| *Mouse_BioGRID* | 0.030 | 0.026 | 0.024 | 0.020 | 0.012 | 0.003 | 0.002 | 0.001 | 0.002 | 0.002 |
| *Worm_BioGRID* | 0.011 | 0.009 | 0.009 | 0.008 | 0.007 | 0.002 | 0.002 | 0.001 | 0.001 | 0.001 |
| *HI_II_14* | 0.042 | 0.038 | 0.033 | 0.032 | 0.028 | 0.006 | 0.008 | 0.006 | 0.003 | 0.003 |
| *Arabidopsis* | 0.107 | 0.095 | 0.092 | 0.088 | 0.079 | 0.015 | 0.018 | 0.010 | 0.008 | 0.007 |
| *Yeast_BioGRID* | 0.048 | 0.045 | 0.037 | 0.037 | 0.029 | 0.024 | 0.023 | 0.015 | 0.014 | 0.013 |
| *Interactome3D* | 0.069 | 0.062 | 0.061 | 0.053 | 0.043 | 0.015 | 0.012 | 0.008 | 0.012 | 0.011 |
| *Lit_BM_13* | 0.024 | 0.022 | 0.026 | 0.023 | 0.017 | 0.006 | 0.005 | 0.004 | 0.004 | 0.004 |
| *Hein* | 0.068 | 0.055 | 0.066 | 0.052 | 0.047 | 0.027 | 0.025 | 0.017 | 0.015 | 0.012 |
| *Fly_BioGRID* | 0.028 | 0.026 | 0.016 | 0.017 | 0.013 | 0.001 | 0.001 | 0.001 | 0 | 0 |
| *Arabidopsis_BioGRID* | 0.116 | 0.106 | 0.098 | 0.096 | 0.088 | 0.015 | 0.016 | 0.006 | 0.008 | 0.007 |
| *HI_III* | 0.045 | 0.039 | 0.038 | 0.036 | 0.034 | 0.013 | 0.014 | 0.011 | 0.005 | 0.005 |
| *Bioplex* | 0.065 | 0.048 | 0.046 | 0.034 | 0.025 | 0.020 | 0.019 | 0.011 | 0.010 | 0.007 |
| *Human_BioGRID* | 0.024 | 0.025 | 0.019 | 0.023 | 0.016 | 0.007 | 0.009 | 0.006 | 0.004 | 0.004 |

**Suppl. Table 4. AUC-PR evaluation of all non-observed interactions for PPI networks.**
For each network 10% of links have been randomly removed (10 repetitions) and the algorithms have been executed in order to assign likelihood scores to the non-observed links in these reduced networks. In order to evaluate the performance, the links are ranked by likelihood scores and the AUC-PR is computed for all the non-observed links in the ranking. The table reports for each network the mean AUC-PR over the random repetitions. The first rows show the mean AUC-PR and the mean ranking over the entire dataset. The networks are sorted by increasing number of nodes $N$.

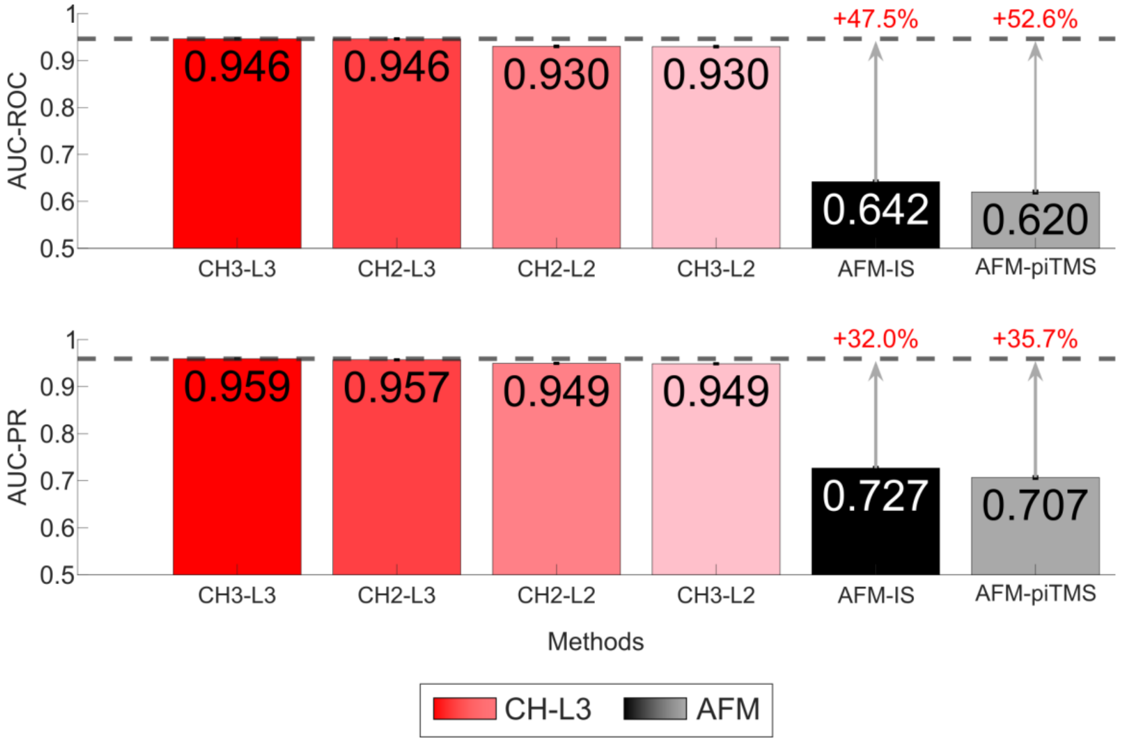


**Suppl. Fig. 5. Comparing the performance of CH and AFM in predicting random 5%GSP+5%GSN protein interactions of the Yeast DIP network.**

Considering CH and AFM methods, the test involves predicting 5% positive and 5% negative interactions in the Yeast DIP network, which are randomly sampled from the GSP (Gold Standard Positives) and GSN (Gold Standard Negatives), respectively. To evaluate the performance, the links are sorted based on the predicted likelihood scores, and AUC-ROC and AUC-PR are computed for the entire sorted list of links. For each evaluation measure, the barplots report the mean performance (with error bars for standard error negligible) of each method on 10 realizations. The color of the CH bars varies depending on the performance value: the higher the performance is, the redder the CH bars appear. The arrows highlight the percentage of improvement between the best CH method and AFM methods as $\frac{CH-AFM}{min(CH,AFM)} \times 100$.

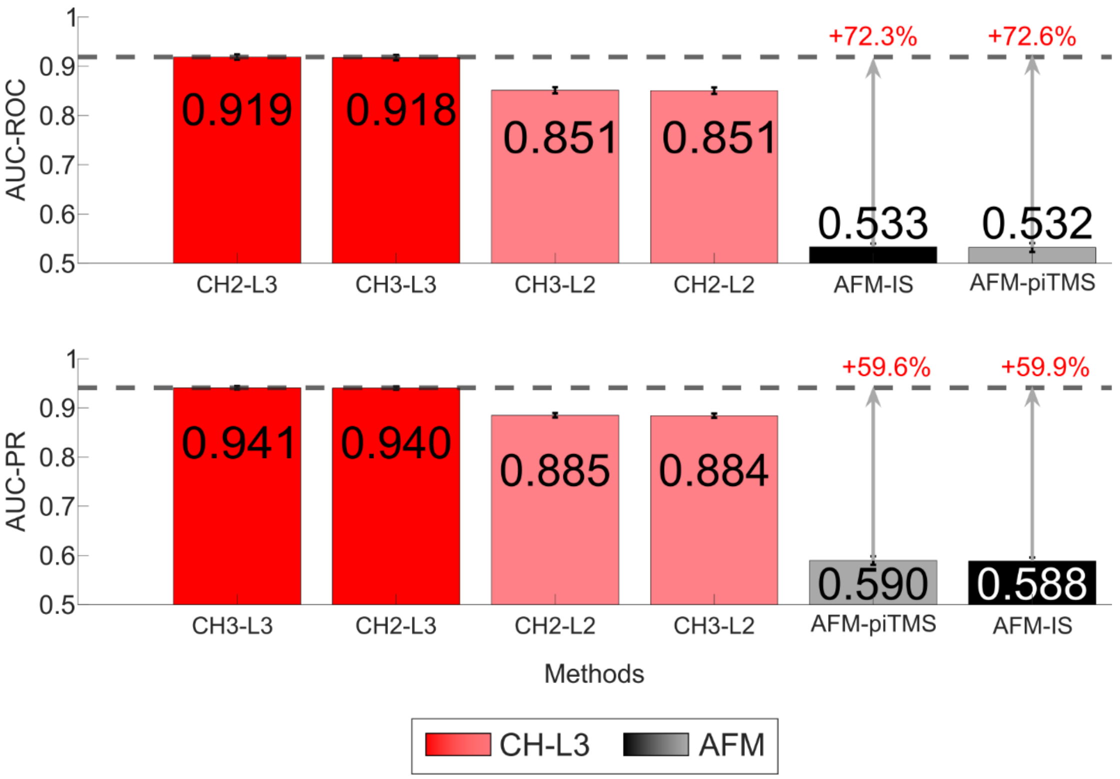


**Suppl. Fig. 6. Comparing the performance of CH and AFM in predicting random 1%positive+1%negative protein interactions of the Yeast DIP network.**

Considering CH and AFM methods, the test involves predicting 1% positive and 1% negative interactions in the Yeast DIP network, which are randomly sampled from the entire positive and negative protein interaction sets, respectively. To evaluate the performance, the links are sorted based on the predicted likelihood scores, and AUC-ROC and AUC-PR are computed for the entire sorted list of links. For each evaluation measure, the barplots report the mean performance (with error bars for standard error which are negligible) of each method on 10 realizations. The color of the CH bars varies depending on the performance value: the higher the performance is, the redder the CH bars appear. The arrows highlight the percentage of improvement between the best CH method and AFM methods as $\frac{CH-AFM}{min(CH,AFM)} \times 100$.

| | *N* | *E* | *k* | *C* | *L* | *LCP-corr* |
|---|---|---|---|---|---|---|
| *S_Pombe_BioGRID* | 1929 | 3276 | 3.40 | 0.06 | 4.67 | 0.66 |
| *Yeast* | 2018 | 2705 | 2.68 | 0.05 | 5.61 | 0.52 |
| *Mouse_BioGRID* | 2099 | 2923 | 2.79 | 0.05 | 6.00 | 0.73 |
| *Worm_BioGRID* | 3153 | 5392 | 3.42 | 0.03 | 4.78 | 0.76 |
| *HI_II_14* | 4392 | 13416 | 6.11 | 0.04 | 4.13 | 0.88 |
| *Arabidopsis* | 4866 | 10928 | 4.49 | 0.10 | 5.18 | 0.56 |
| *Yeast_BioGRID* | 4888 | 27358 | 11.19 | 0.12 | 3.60 | 0.85 |
| *Interactome3D* | 5291 | 6034 | 2.28 | 0.10 | 7.68 | 0.72 |
| *Lit_BM_13* | 5545 | 10155 | 3.66 | 0.06 | 5.31 | 0.74 |
| *Hein* | 6150 | 28309 | 9.21 | 0.15 | 3.90 | 0.92 |
| *Fly_BioGRID* | 7283 | 23751 | 6.52 | 0.01 | 4.33 | 0.74 |
| *Arabidopsis_BioGRID* | 7622 | 28547 | 7.49 | 0.10 | 4.81 | 0.61 |
| *HI_III* | 7854 | 43939 | 11.19 | 0.06 | 3.83 | 0.86 |
| *Bioplex* | 10961 | 56553 | 10.32 | 0.10 | 4.28 | 0.88 |
| *Human_BioGRID* | 12605 | 65025 | 10.32 | 0.07 | 3.65 | 0.79 |
| *Human_Menche* | 13460 | 138427 | 20.57 | 0.17 | 3.58 | 0.97 |

**Suppl. Table 5. Statistics for PPI networks.**

The table reports for each network several statistics: $N$ is the number of nodes; $E$ is the number of edges; $k$ is the average node degree; $C$ is the average clustering coefficient, computed for each node as the number of links between its neighbours over the number of possible links; $L$ is the characteristic path length of the network; *LCP-corr* is the Local-Community-Paradigm correlation [4], representing the correlation between the number of common-neighbours and the number of links between them, looking at each pair of connected nodes in the network. The networks are sorted by increasing $N$.

| | *N* | *E* | *k* | *C* | *L* | *LCP-corr* |
|---|---|---|---|---|---|---|
| *Yeast_DIP* | 4951 | 22382 | 9.04 | 0.10 | 3.95 | 0.81 |

**Suppl. Table 6. Statistics for the Yeast DIP network used in CH-AFM comparison.**

The table reports several statistics for the Yeast DIP network: $N$ is the number of nodes; $E$ is the number of edges; $k$ is the average node degree; $C$ is the average clustering coefficient, computed for each node as the number of links between its neighbours over the number of possible links; $L$ is the characteristic path length of the network; *LCP-corr* is the Local-Community-Paradigm correlation [4], representing the correlation between the number of common-neighbours and the number of links between them, looking at each pair of connected nodes in the network.

| Network | Species | Source/Reference |
|---|---|---|
| *Arabidopsis* | *A. Thaliana* | [5] |
| *Arabidopsis_BioGRID* | *A. Thaliana* | BioGRID v. 3.4.143 [6] |
| *Worm_BioGRID* | *C. Elegans* | BioGRID v. 3.4.143 [6] |
| *Fly_BioGRID* | *D. Melanogaster* | BioGRID v. 3.4.143 [6] |
| *HI_II_14* | *H. Sapiens* | [7] |
| *Interactome3D* | *H. Sapiens* | [8] |
| *Lit_BM_13* | *H. Sapiens* | [7] |
| *Hein* | *H. Sapiens* | [9] |
| *HI_III* | *H. Sapiens* | unpublished (link) |
| *Bioplex* | *H. Sapiens* | [10] |
| *Human_BioGRID* | *H. Sapiens* | BioGRID v. 3.4.143 [6] |
| *Human_Menche* | *H. Sapiens* | [11] |
| *Mouse_BioGRID* | *M. Musculus* | BioGRID v. 3.4.143 [6] |
| *Yeast* | *S. Cerevisiae* | [12] |
| *Yeast_BioGRID* | *S. Cerevisiae* | BioGRID v. 3.4.143 [6] |
| *Yeast_DIP* | *S. Cerevisiae* | DIP database |
| *S_Pombe_BioGRID* | *S. Pombe* | BioGRID v. 3.4.143 [6] |

**Suppl. Table 7. Sources for PPI networks.**

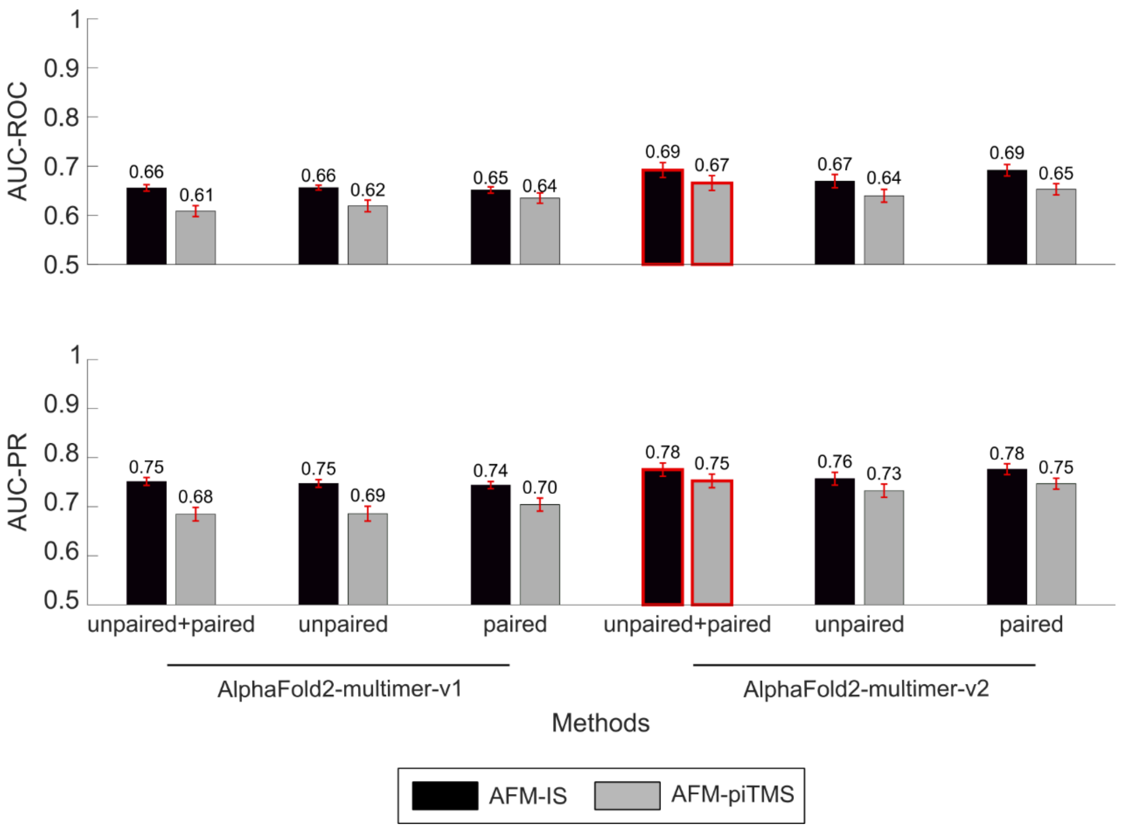

**Suppl. Fig. 7. Determining AFM settings with highest performance in predicting the GSP and GSN protein pairs of the Yeast DIP network.**

For the Yeast DIP network and for each combination of the pair-mode (“unpaired+paired", "unpaired", "paired") and the model-type ("AlphaFold2-multimer-v1", "AlphaFold2-multimer-v2"), the evaluation of the structural-based interaction predictions is reported. The AUC-ROC and the AUC-PR are computed at each 1% increment of the entire GSP (4476 interactions) and the entire GSN (4476 interactions) respectively. Note that AFM sporadically failed to successfully predict some interactions, therefore the final number of predicted GSP and GSN marginally vary of few units across the different settings. The bar plots show these mean performances, as well as the standard errors of the mean. The mean performance is displayed on top of the error bars. The best AFM setting is highlighted in red.
